# ^19^F Ultrafast MAS NMR Reveals the Dynamic Basis of pH-Dependent Regulation in Proteorhodopsin

**DOI:** 10.64898/2026.07.18.739351

**Authors:** Samuel Seidl, Danhua Dai, Anouk Ebenezer, Jennifer Anne Winzer, Johanna Becker-Baldus, Xiao He, Clemens Glaubitz

## Abstract

^19^F NMR spectroscopy is a powerful approach for studying complex biomolecular systems because of its high sensitivity, exceptional responsiveness to local structural changes, and simplified spectra. Here, we demonstrate the application of ^19^F ultrafast MAS NMR to 5-fluorotryptophan-labelled proteorhodopsin reconstituted in lipid bilayers. By assigning 9 of the 10 tryptophan resonances, pH-dependent analyses of chemical shifts, line shapes, and conformational exchange reveal the dynamics of two functionally important residues: W34 in the interprotomer His-Asp-Trp triad and W98 within the retinal-binding pocket. The results identify W34 as a dynamic regulator of proton transport and support a model in which slow ring flipping on the seconds timescale transiently modulates the W34-H75 interaction, thereby acting as a pH-dependent molecular throttle. The spectral characteristics of W98 further suggest that it functions as a dynamic regulator of the photocycle within the retinal-binding pocket. Beyond these mechanistic insights, we show that a MAS rate of 100 kHz markedly enhances the resolution of this ^19^F-labelled membrane protein. Combined with a simple chemical-shift scoring metric and advanced, linear-scaling AF-QM/MM-based ^19^F chemical shift calculations of all sites within this protein, this workflow provides a robust and broadly applicable framework for characterizing membrane protein structure and dynamics in native-like lipid environments.

**TABLE OF CONTENT GRAPHICS:** 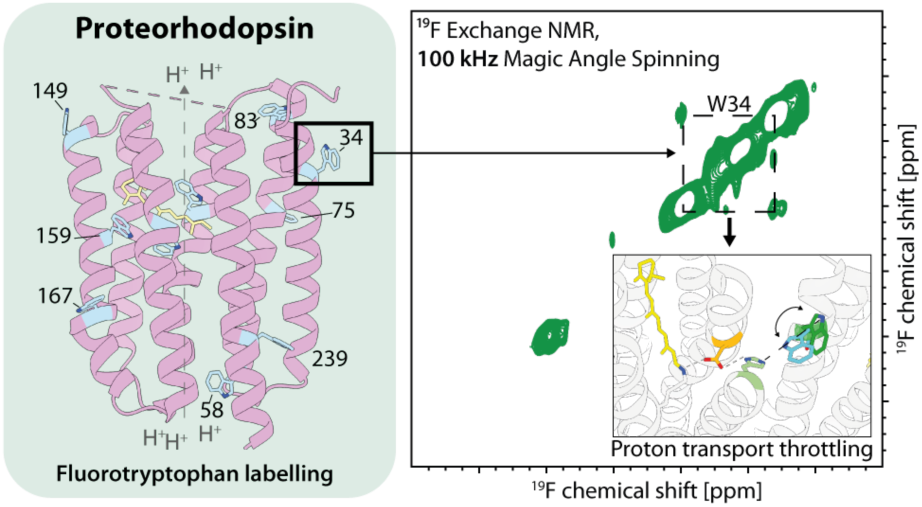

## INTRODUCTION

Tremendous progress in the structure determination of membrane proteins - the interface between the cell and its surroundings – has been achieved over the past decades through X-ray crystallography and cryo-electron microscopy. Visualizing the architecture of these often challenging systems is a crucial first step toward understanding their roles in energy generation, transport and signal transmission across biological membranes. However, even with advances in time-resolved structural studies, the conformational equilibria and the shifts within them that underlie protein function and malfunction remain largely elusive.

NMR spectroscopy has played a pivotal role in elucidating the relationship between protein conformation and function by providing direct access to protein dynamics and conformational landscapes.^1–5^ However, extending these studies to large membrane proteins remains challenging. In solution-state NMR, slow molecular tumbling leads to severe line broadening, whereas in solid-state NMR, uniformly labeled samples often suffer from extensive spectral overlap. Site- or residue-selective labelling strategies can alleviate these limitations. In particular, the incorporation of ^19^F offers several unique advantages: Its high gyromagnetic ratio and 100% natural abundance provide great detection sensitivity, while its wide chemical shift range and pronounced sensitivity to the local electrostatic environment make it an excellent structural reporter.^6^ Consequently, the resulting NMR spectra are relatively simple with a reduced number of signals. These features made ^19^F NMR a powerful approach for probing the conformational landscape of large biomolecular systems in solution,^7–8^ establishing fluorine nuclei as highly sensitive conformational reporters. However, when considering large membrane proteins in native-like environments such as liposomes, solution-state NMR becomes impractical. Under these conditions, strong dipolar couplings and especially relaxation due to the large chemical shift anisotropy (CSA) of some ^19^F-labels, which increases with the square of the magnetic field, results in severe line broadening.^4^

Recent advances in solid-state NMR have focused on implementing faster magic-angle spinning (MAS) rates of 100 kHz and above to enable direct proton detection. ^9–10^ Thereby the strong homonuclear dipolar couplings are averaged enabling high-resolution spectra and side-chain assignments even in large protein systems.^11^ Fast MAS requires the use of smaller rotors. The smaller sample amount is to some extent compensated for by the vicinity of the coil to the sample, the higher sensitivity of protons and thus experiments on a small amount of sample become feasible which is of great importance when studying biological systems.

Fast MAS NMR has also been shown to be beneficial for studying ^19^F-labelled samples as the strong heteronuclear ^19^F-^1^H dipole-dipole couplings as well as the large ^19^F CSA can be effectively removed. Several recent studies explored a combination of fast MAS at 60 kHz with ^19^F NMR in the context of distance and structure determination for viral and membrane proteins.^12–14^ Spinning at 111 kHz MAS with ^19^F detection was used for the characterization of active pharmaceutical compounds demonstrating a progressively improving resolution with increasing MAS rate.^15^ In case of membrane proteins, a recent study from our lab demonstrated detection of a fluorinated ligand bound to a G protein-coupled receptor (GPCR) under 100 kHz MAS NMR conditions.^16^ Other previous reports on chemically labelled proteorhodopsin^17^, A2A adenosine receptor^18^ and diacylglacerolkinase using unnatural amino acids for labelling^19^ were carried out using MAS rates between 10 and 40 kHz.

Here, we explore the use of ultrafast 100 kHz MAS for a ^19^F NMR study on fluorinated tryptophans in the integral membrane protein proteorhodopsin in its lipid environment. Proteorhodopsins (PRs) are light-driven outward proton pumps found in marine bacteria contributing to their metabolism by phototrophy. Probably, they are the most abundant retinal-based proteins on Earth.^20^ Their abundance in planktonic microorganisms imply an important contribution to bioenergetics and ocean ecology.^21^ PR is highly adapted to the environmental conditions with a green-light absorbing form found in bacteria in shallow waters and a blue-light absorbing form in deeper sea levels.^22^

For our study, green light-absorbing proteorhodopsin (GPR) has been chosen which is the first discovered PR and was found in the SAR86 clade in marine γ-proteobacteria.^23^ It consists of a heptahelical transmembrane bundle (Fig. 1) which is typical for rhodopsins but also a structural feature which is shared with GPCRs. In the membrane, it assembles mainly into pentamers. GPR contains retinal as a prosthetic group which is bound by a Schiff base to a conserved K231. The retinal is sandwiched between W197 on the intracellular and W98, close to the Schiff base, on the extracellular side (Fig. 1). Another tryptophan, W159 on the intracellular side is found in proximity to the β-ionone ring. All three are highly conserved within the PR family and W98 and W197 are also found in archaeal rhodopsins e.g., the prototypical archaeal bacteriorhodopsin from *H. salinarum* and eukaryotic rhodopsins. W159 and W197 contribute to the green/blue color switching mechanism.^24^

**Figure 1:**
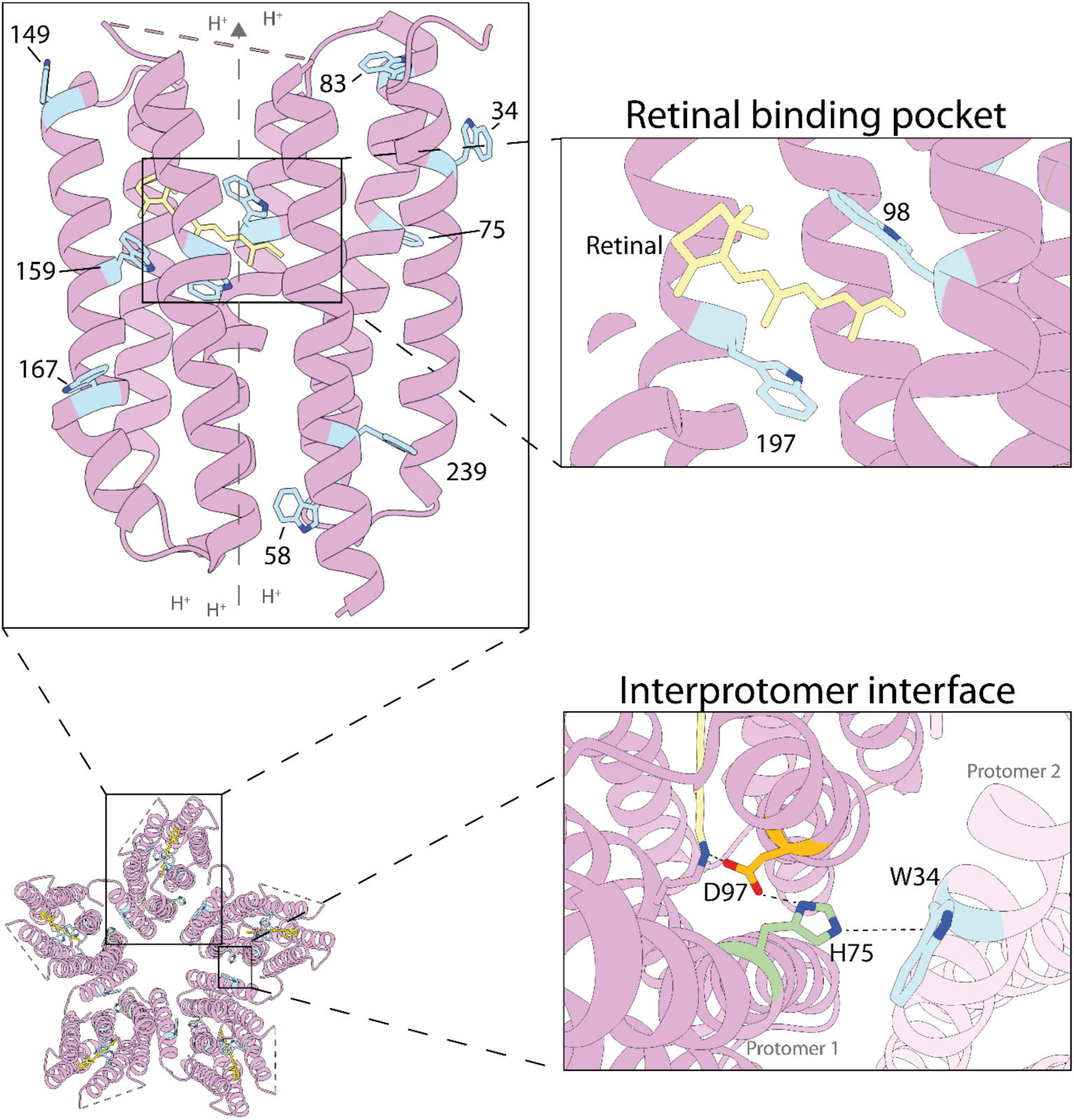
Structural overview of green light-absorbing proteorhodopsin. Cryo-EM structure (PDB ID 7b03) of GPR shows a pentameric assembly. The protomer consists of 10 tryptophans (blue) with several being functionally or structurally relevant. W98 and W197 are part of the retinal (yellow) binding pocket and conserved across most rhodopsins. W34 residues form interprotomeric contacts to the neighboring H75-D97 cluster residues (green) modulating the transport in the neighboring monomer.

Proton pumping in proteorhodopsin is initiated by light absorption through the retinal triggering an isomerization from all-*trans*-retinal to 13-*cis*-retinal which is the first step of the photocycle. The photocycle describes a number of protonation and deprotonation events along with structural changes within the protein. Some of these structural changes are transduced from the deformed retinal by W197 to the protein as seen for bacteriorhodopsin.^25^ Despite its conserved nature, a similar role has not been described for W98. After isomerization, the conserved proton acceptor D97, exhibiting an unusually high pK_a_ value for an aspartate, is protonated by the Schiff base. In the final stages of the photocycle, the proton is further translocated to a proton release group so that it is released into the extracellular media and re-isomerization of the retinal occurs in order to return to the ground state. The unusually high pK_a_ of D97 is caused by hydrogen bonding to H75.^26^ Together they form the conserved His-Asp cluster. Previous solid-state NMR studies have shown that His75 performs a ring flipping motion during the photocycle thus controlling the acceptance and release of protons by D97.^27^

Additionally, recent pH-dependent structural snapshots of marine actinobacter proteorhodopsin have shown that under low pH conditions, which reduces proton transport, the histidine is hydrogen bonded to the aspartate in contrast to high pH conditions, where the imidazole ring is rotated.^28^ In γ-proteobacteria, an additional layer of modulation has been suggested in form of interprotomeric contacts of the conserved W34 of one protomer to the His-Asp cluster in the neighboring protomer as shown by mutagenesis and solid-state NMR.^28–29^. Due to the high pK_a_ value of D97, this His-Asp-Trp triad might act as a pH sensor.^27^

Given the strategic positions of tryptophan residues in GPR and their proposed functional roles, we employed 5-fluorotryptophan as a probe to characterize their pH-dependent dynamic landscape and their contribution to pH-dependent transport regulation. We assigned the corresponding ^19^F resonances through site-directed mutagenesis and rationalized the observed chemical shift dispersion using local structural features together with chemical shift calculations based on an automated fragmentation quantum mechanics / molecular mechanics (AF-QM/MM) approach. Predicting these shifts in the context of a large membrane protein requires methods that explicitly account for the surrounding protein and solvent environment. The AF-QM/MM method enables accurate and efficient linear-scaling *ab initio* quantum chemical calculations of NMR parameters of large proteins. This method has been previously demonstrated successfully in calculating ^15^N and ^13^C chemical shifts in other membrane proteins and PR^24, 30^ and is now applied to the more challenging task of predicting ^19^F chemical shifts. The analysis of pH-dependent conformational exchange and NMR line shapes revealed distinct dynamic signatures and led us to propose a motion-based, interprotomeric mechanism of pH-dependent regulation mediated by the conserved His–Asp–Trp cluster in proteorhodopsins.

## RESULTS

### Biochemical characterization of [5-^19^F-Trp]-wtGPR

Incorporation of 5-fluorotryptophan was achieved by adding it to the culture medium during expression. SDS-PAGE and BlueNative-PAGE of [5-^19^F-Trp]-wtGPR after metal affinity purification showed identical running behavior to unlabeled protein (Fig. S1A, B). An additional size-exclusion chromatography step (SEC) was introduced to remove non-functional aggregates, which occur in [5-^19^F-Trp]-wtGPR preparations (Fig. S1C).

To evaluate whether the fluorinated amino acid changed the optical properties of GPR, UV-Vis spectra of labelled and the unlabeled protein were compared. At pH 7.5, both show the same absorption maximum of the retinal chromophore at λ_max_ = 520 nm (Fig. 2A, inset). A pH titration of λ_max_, which reflects the pKa of the primary proton acceptor D97, reveals a shift to lower values upon 5-^19^F-Trp labelling (Fig. 2A). Additionally, the pH-dependent wavelength dispersion is narrower with an upper limit at 536 nm in contrast to 546 nm in the unlabeled protein.

**Figure 2:**
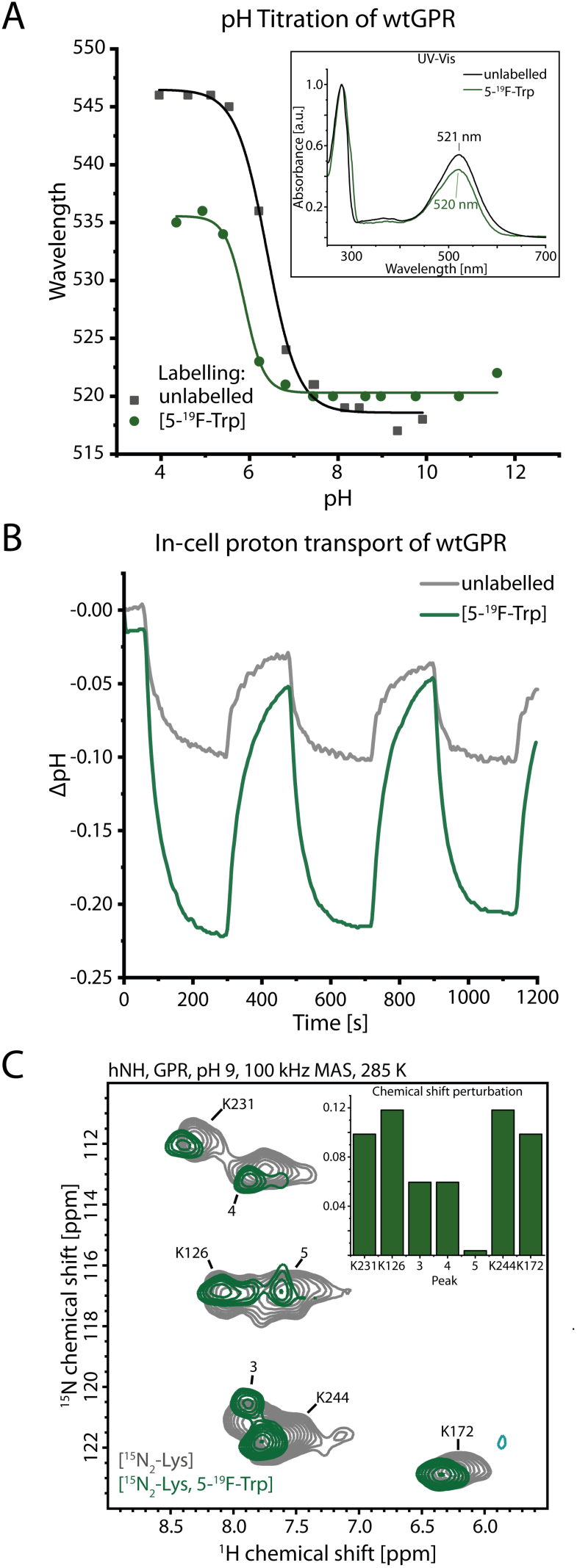
Biochemical characterization of [5-^19^F-Trp]-wtPR. **(A)** pH titration of unlabeled GPR and [5-^19^F-Trp]-wtGPR. The absorption of the bound retinal was monitored during pH titration and the obtained points were fitted with a sigmoidal function. The pK_a_ for GPR was 6.4 ± 0.1 with a wavelength dispersion between 519 nm – 546 nm and for labelled GPR 5.9 ± 0.1 with a wavelength dispersion between 520 nm – 536 nm. The inset shows the UV-Vis spectra of labelled and unlabeled wtGPR at pH 7.5. **(B)** In-cell proton transport unlabeled (black) and labelled GPR (green) expressed in *E. coli* cells. The pH was monitored over time. For certain periods of time, the cell suspension was illuminated with green light (520 nm) during the measurement. **(C)** hNH at 100 kHz MAS of [^15^N_2_-Lys]-wtGPR (black) and [^15^N_2_-Lys, 5-^19^F-Trp]-wtGPR (green) reconstituted in liposomes. Lysine residue assignments from previous assignments^11^ are indicated. The insert shows the small chemical shift perturbations of the lysines between the two preparations.

The effect of fluorination upon proton pumping was directly probed by an in-cell proton transport assay in *E. coli* cells overexpressing GPR. Upon illumination with green light, GPR starts proton pumping into the extracellular medium, thus lowering the pH values. After illumination, the pH of the medium increases again due to influx of protons into the cells. Transport was observed in the unlabeled as well as in the ^19^F-labelled GPR showing a functionally intact protein (Fig. 2B). Surprisingly, [5-^19^F-Trp]-GPR expressing cells show a larger pH change compared to those expressing unlabeled GPR.

After purification the protein was reconstituted in DMPC/DMPA liposomes, sedimented in 0.7 mm MAS rotos and global integrity of the protein was probed by proton-detected ssNMR hNH spectra at 100 kHz MAS. Therefore, the protein was additionally labelled with ^15^N_2_-lysine which results in labelling of the residue which is linked to the chromophore as well as six additional residues (Fig. S2). The hNH spectrum on [^15^N_2_-Lys]-wtGPR shows seven well separated backbone resonances (Fig. 2C). All of these signals can also be observed in the spectrum of [^15^N-Lys, 5-^19^F-Trp]-GPR showing very similar chemical shifts and thus proving the structural integrity of the protein.

### MAS- and temperature-dependent spectra

Fast MAS is needed to average the ^1^H-^19^F dipolar couplings. To analyze to which extend the MAS frequency influences the spectra, ^19^F MAS-dependent spectra with MAS rates between 40 and 100 kHz were recorded (Fig. 3A) and show improved signal-to-noise ratio and resolution with increasing rotation frequency. In addition, we observed a temperature dependence on the spectral resolution (Fig. 3B bottom) showing a significant improvement in resolution at the higher temperature due to the increased protein dynamics. We performed a sample temperature scan (Fig. S3) and the optimal resolution was reached at 298 K which was the highest temperature tested. This temperature was then chosen for all subsequent experiments.

**Figure 3:**
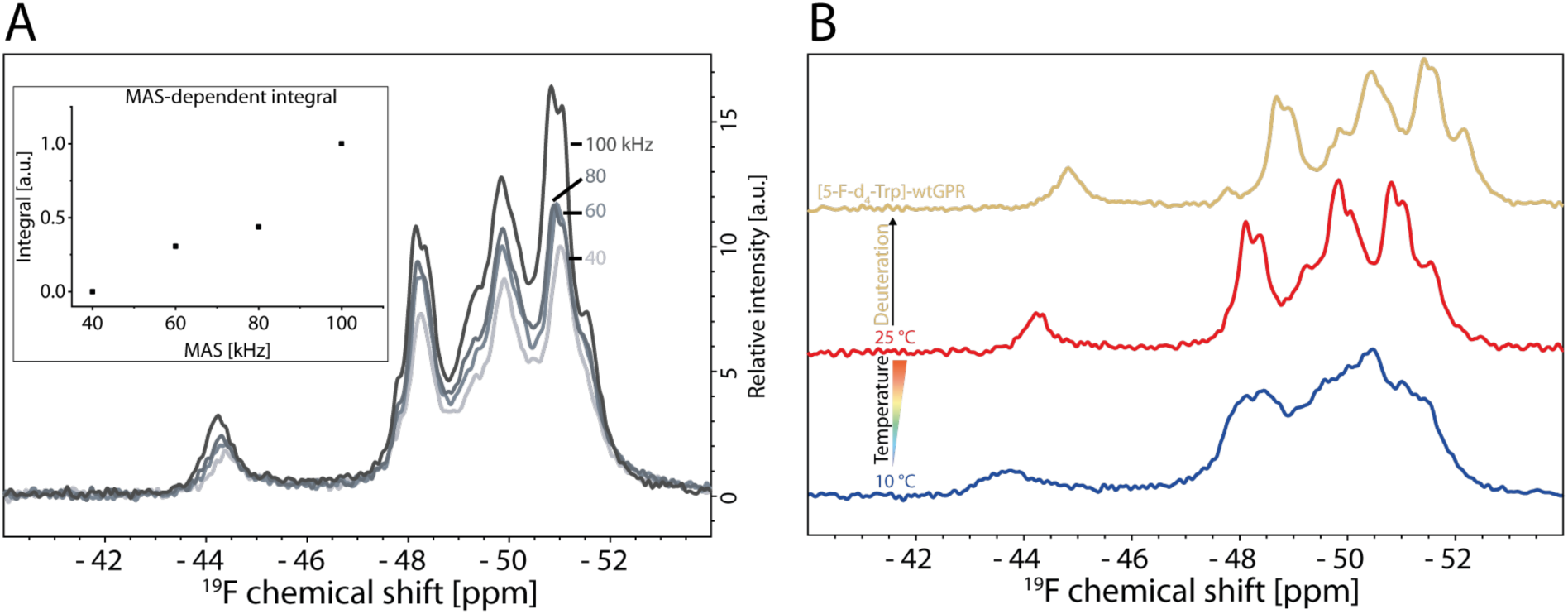
^19^F-MAS NMR spectra of [5-^19^F-Trp]-wtGPR. **(A)** MAS-dependent spectra of [5-^19^F-Trp]-wtGPR. ^19^F spectra were recorded at 40, 60, 80 and 100 kHz MAS. In order to compensate for different sample temperatures due to the MAS, cooling was adjusted in order to have the same chemical shift of water in the ^1^H detected spectra. The insert shows the integral of the whole spectrum in dependence of the MAS rate. **(B)** Linewidth contributions in ^19^F-detected ultra-fast MAS measurements. [5-^19^F-Trp]-GPR in liposomes was recorded at two different sample temperatures at 100 kHz MAS (Red and blue spectrum). The sample temperature was estimated using the temperature dependent proton chemical shift of water. The yellow spectrum was recorded with [5-^19^F-d_4_-Trp]-wtGPR at 25 °C.

To probe whether further improvements can be achieved by disrupting the local proton coupling network and hence suppressing residual ^1^H-^19^F dipole-dipole couplings, deuterated 5-^19^F-d_4_-Trp was incorporated. The [5-^19^F-d_4_-Trp]-wtGPR spectrum exhibits a high-field shift of approximately 0.55 ppm (Fig. 3B top), due to the isotope effect.^31^ However, linewidths obtained by deconvolution of the spectrum (as described below) showed no significant improvements in resolution (Figs. 4A, S4; Tables S1, S2) indicating that the residual dipolar contributions to the linewidth at this spinning rate is minor. Therefore, all further samples in this work were prepared using protonated 5-^19^F-Trp.

**Figure 4:**
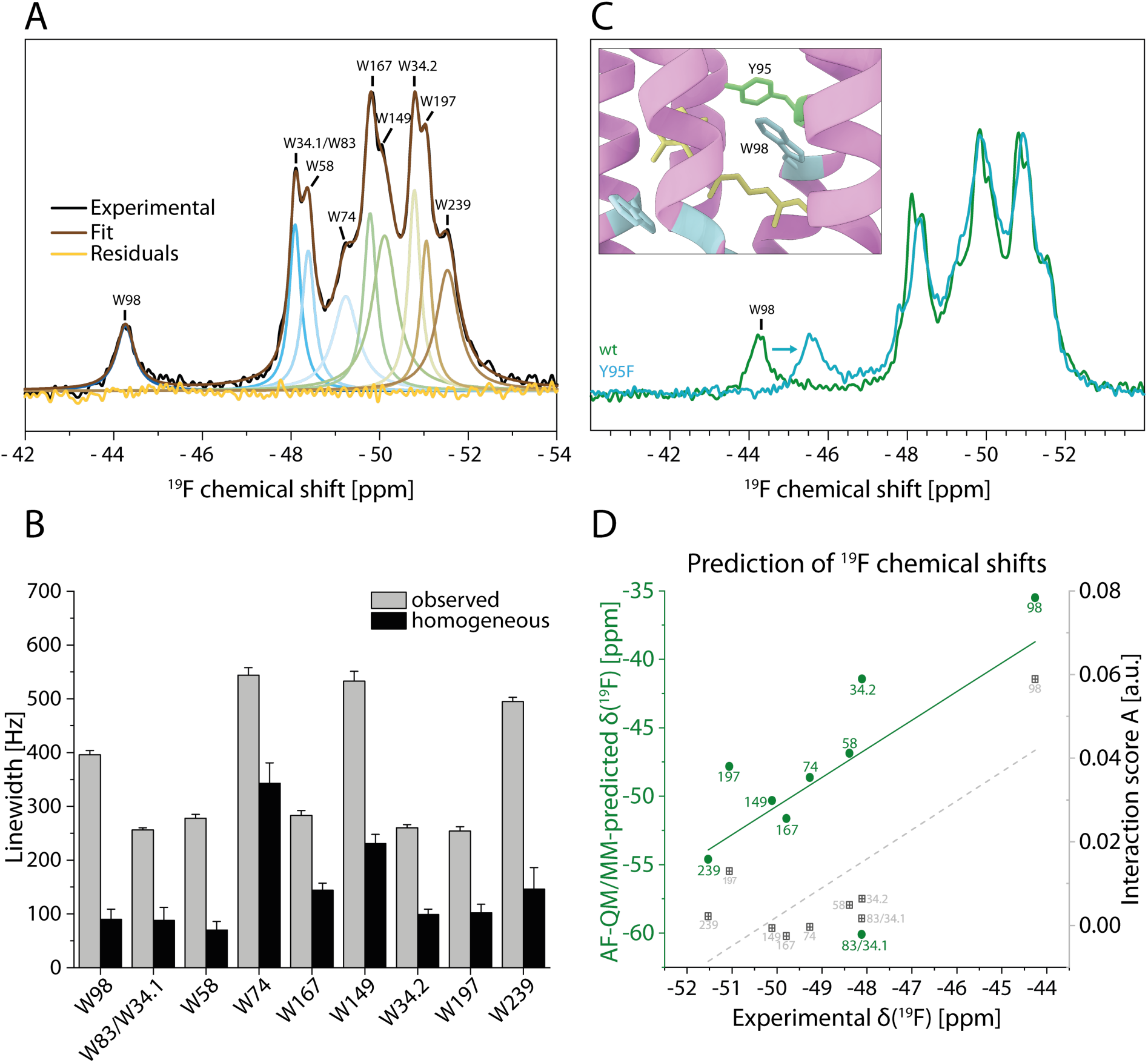
Site-specific characterization of wtGPR. **(A)** Summary of 5-^19^F-Trp assignment in [5-^19^F-Trp]-wtGPR and Lorentzian deconvolution of the spectrum (top panel). A nine-state Lorentzian fit was performed according to the nine distinguishable signals. Additionally, the sum of the deconvolution is overlayed with the spectrum and the residuals are shown. **(B)** T_2_-determined, homogeneous and observed linewidth for 5-^19^F-Trp residues in [5-^19^F-Trp]-wtGPR. Error bars represent the numerical errors obtained by the fitting procedure. **(C)** Exemplary assignment of 5-^19^F-Trp residues in GPR on the basis of [5-^19^F-Trp]-GPR-Y95F. **(D)** ^19^F chemical shift prediction of fluorotryptophans in [5-^19^F-Trp]-wtGPR. The absolute predictions based on AF-QM/MM calculations are shown in green with the numbers indicating the corresponding residue. A linear regression was performed between the calculated and experimental chemical shifts (solid green line). An interaction score based on the cryo-EM structure (PDB ID: 7b03) of GPR was determined (grey). The dispersion of the scores was also correlated to the experimental chemical shifts by linear regression (dashed grey line), see also Table S3.

### Chemical shift and line shape analysis

In the ^19^F spectrum recorded at 100 kHz and 298 K, nine signals from 10 tryptophan residues can be distinguished. For further analysis, the spectrum was deconvoluted assuming Lorentzian line shapes (Fig. 4A) and assigned, see below. The obtained peaks differ significantly in their respective line width from each other ranging from 250 to 540 Hz (Table S2).

In order to assess whether the observed line widths are dominated by inhomogeneous (e.g., heterogenous distribution of conformations) or homogeneous (e.g., conformational dynamics, insufficient ^1^H-^19^F decoupling) contributions, Hahn echo experiments were conducted to determine individual T_2_ relaxation times and, consequently, the homogeneous line widths (Figs. 4B, S5, Table S2). The resulting homogeneous line widths range from 70 to 340 Hz and are smaller than the corresponding values obtained by spectral deconvolution for all sites. These findings show that the observed spectra contain both homogeneous and inhomogeneous broadening contributions. The inhomogeneous component may arise from conformational dynamics that are slow on the NMR time scale or from the coexistence of multiple conformations with distinct chemical shifts. The data show that the ^19^F-labelled sites not only differ in their chemical shifts but also in line width contributions, revealing site-specific differences in conformation and dynamics.

To assign the signals to the ten tryptophans that occur in GPR, a mutagenesis approach was applied (Figs. 4A, S6, S7). 19F-Trp-GPR samples with single tryptophan to phenylalanine mutation could be prepared for eight out of ten residues (Fig. S6). However, expression of constructs with W98F or W197F failed probably due to their critical structural role in GPR. Both residues are part of the retinal binding pocket. Therefore, “nudge” mutations, in which the structurally nearest residues to the target site were mutated, were used for assignment. Figure 4C shows the ^19^F spectrum of the Y95F “nudge” mutation, which enabled the assignment of the nearby W98 residue through induced chemical shift perturbation of its signal. This combined approach enabled the assignment of all resolved signals. Eight resonances were assigned to specific Trp residues, whereas W159 could not be assigned. W34, the tryptophan located at the protomer interface, exhibited two resonances, one of which overlapped with the W83 signal (see SI for further details).

An interesting observation is the unequal dispersion of the 5-^19^F-Trp signals between -44 and -52 ppm. The W98 signal is isolated and experiences the highest downfield shift whereas the other signals partially overlap. Solvent exposure could be one factor influencing the observed dispersion.^32–33^ In order to assess the influence of solvent contacts, an MD simulation of the cryo-EM structure (PDB ID: 7b03) within lipid bilayers from the MemProtMD database was consulted.^34^ To investigate this relationship, the solvent contact scores derived from the simulations were correlated with the experimentally determined chemical shifts (Fig. S9A, Table S3). However, no significant correlation was observed between solvent contact scores and chemical shifts.

We then tried to correlate ^19^F chemical shifts, which should be sensitive to the electrostatic and van-der-Waals environments, with structural properties.^6, 35–36^ A simplified approach considering previous publications^35–36^ was used, were a simple, empirical score A (Eq. 1) based on charge (+2), polar (+1) or an unpolar (-1) interaction for all atoms within a 10 Å radius around the 5-^19^F-Trp (determined by ChimeraX^37^), was defined. This interaction was weighted by the distance between the corresponding atom and the fluorine label. A charge-dipole interaction was weighted with 1/r^4^ and dipole-dipole (as Keesom interaction) or van-der-Waals interaction was weighted with 1/r^6^. The interaction score A is then defined as:

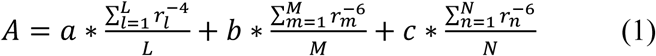

where L, M and N are the number of atoms within the considered volume for each interaction type and a = +2 Å^4^, b = +1 Å^6^, c = -1 Å^6^.

All contacts to the retinal moiety are treated as charged interaction because of the delocalized charge of the Schiff base. With this very simple and empirical approach, a reasonable linear correlation between the obtained scores and the experimental chemical shifts is obtained (Fig. 4D, grey points). Especially, the chemical shift separation between W98 and the other residues is well reflected in these scores.

To provide further quantitative support for the observed ^19^F chemical shifts, we performed absolute ^19^F chemical shift predictions based on the GPR cryo-EM (PDB 7b03)^38^ structure using an AF-QM/MM approach.^39^ The AF-QM/MM method enables accurate and efficient linear-scaling ab initio quantum chemical calculations of NMR parameters of large proteins. This method has been previously demonstrated successfully in calculating ^15^N and ^13^C chemical shifts in other membrane proteins and PR^24, 30^ and is now applied to the more challenging task of predicting ^19^F chemical shifts.^40–41^ Following validation of the protocol with a 5 Å cutoff distance (see SI, Fig. S10 and Fig. S11 for parametric evaluation of cutoff distance and ion effects), we systematically tested two aspects: (A) structural model, comparing monomer-based predictions with pentamer-based predictions where the chemical shift was averaged over the five protomers; and (B) treatment of long-range electrostatics, comparing isolated QM calculations of the core and buffer regions with QM/MM calculations incorporating environmental point charges. Among the four combinations, the monomer model incorporating environmental charges performed most consistently, reproducing the overall experimental chemical shift dispersion with an accuracy comparable to the structure-based classification (Fig. 4D). While the prediction for W83 deviated notably from experiment, the overall agreement supports the spectral assignment and provides independent validation of the observed chemical shift trends. When excluding the outlier W83, the monomer-based AF-QM/MM predictions outperformed the structure-based classification (Table S3). The beneficial effect of including environmental charges was further confirmed by systematic comparison with isolated QM calculations (Table S3, Fig. S9B).

### pH-dependent conformational changes in GPR

As GPR functions as a light-driven proton pump, varying the pH can be used to mimic different functional states of the protein,^29, 38^ particularly by modulating the protonation state of the primary proton acceptor, D97. Fig. 5A shows ^19^F spectra of [5-^19^F-Trp]-GPR recorded at pH values below (pH 5), near (pH 6) and above (pH 7.5 and 9) the pK_a_ of D97. A general trend is a progressive loss of spectral resolution with decreasing pH. Most notably, the W98 resonance broadens dramatically and is nearly undetectable at pH 5. To exclude irreversible structural destabilization at low pH, proteoliposomes equilibrated at pH 5 (red spectrum) were subsequently exchanged into buffer at pH 7.5 (blue spectrum). This treatment partially restored the spectral resolution, yielding narrower linewidths than those observed at pH 6, demonstrating that exposure to pH 5 does not irreversibly perturb the protein. Consistent with this conclusion, lysine fingerprint spectra recorded at pH 5 confirm that the overall protein fold remains intact (Fig. S12).

**Figure 5:**
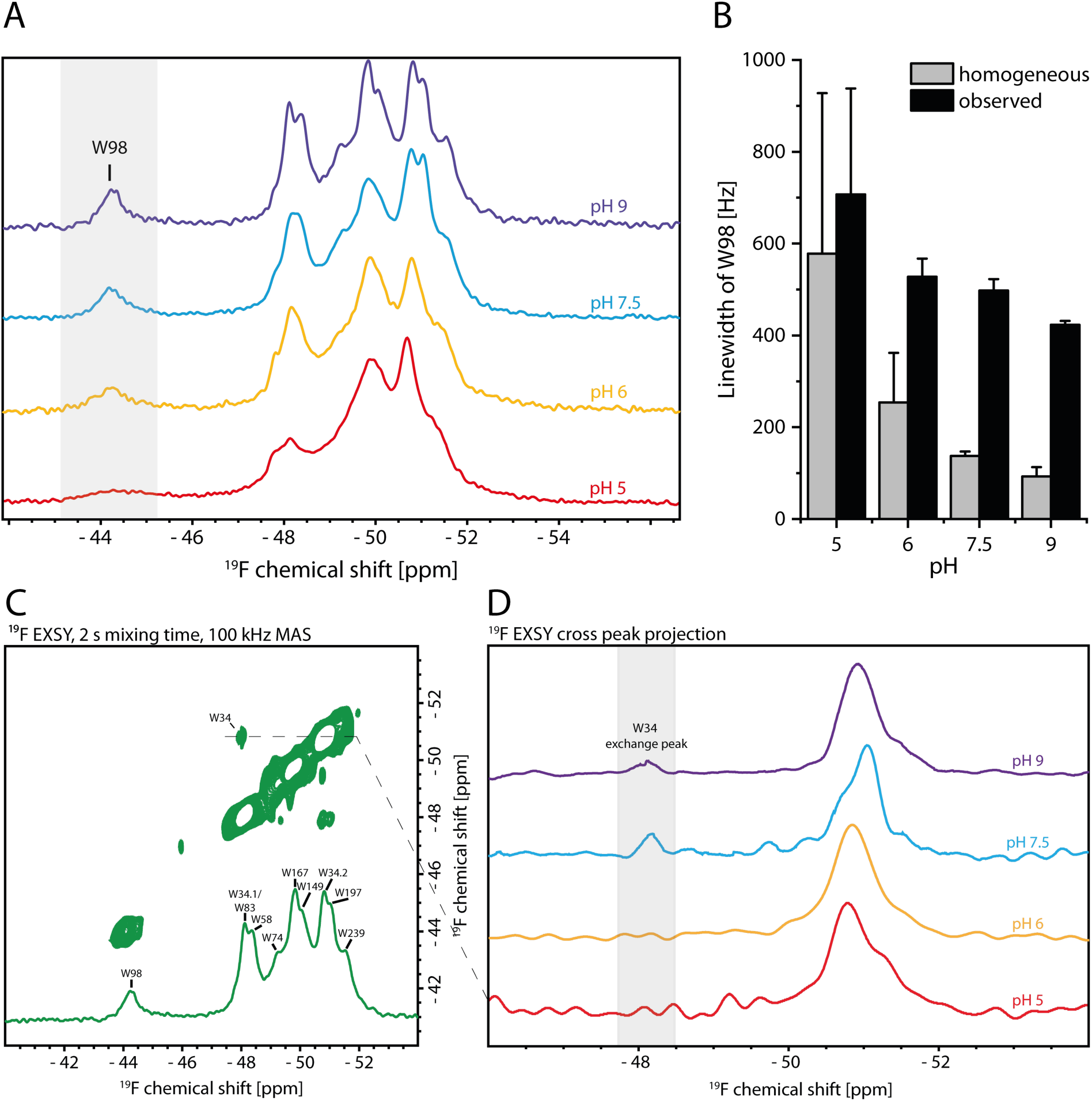
pH-dependent response of tryptophans in [5-^19^F-Trp]-wtGPR. **(A)** pH dependent ^19^F spectra of [5-^19^F-Trp]-wtGPR recorded at different pH values. For pH 7.5 the liposomes which were at pH 5 (red spectrum) were washed with the corresponding buffer to reach pH 7.5. The changes in W98 are highlighted (grey bar) for subsequent analysis. **(B)** pH-dependent observed and refocused linewidths of W98. Linewidths were obtained by deconvolution of the corresponding spectrum or corresponding Hahn echo series of experiments. Errors indicate the fitting error. **(C)** ^19^F EXSY spectra of [5-^19^F-Trp]-wtGPR at pH 9 were recorded at 100 kHz MAS. The EXSY was overlayed with the 1D ^19^F Hahn echo spectrum and the corresponding assignment. **(D)** pH-dependent exchange of W34. The cross-peak volume was normalized to the signal of the diagonal in pH-dependent EXSY spectra of [5-^19^F-Trp]-wtGPR spectra.

To analyze the nature of the observed line broadening, Hahn Echo T_2_ measurements were carried out. Due to the increased line broadening at lower pH values, we restrict the analysis to the resolved signal of W98. The homogeneous linewidth as well as the observed linewidth decrease with increasing pH (Fig. 5B).

To probe slow motions, 2D exchange (EXSY) spectra were recorded at pH 9 (Figs. 5C, S13). A cross peak at -50.7 ppm / -47.9 ppm was observed. Such a cross peak can either result from slow exchange between two states which differ in chemical shift or from spin diffusion between ^19^F atoms which are close in space. Based on the cryo-EM structure (PDB 7b03), we analyzed all the possible ^19^F-^19^F distances in the sample. The distances between the ^19^F nuclei in W159 and W197 within the retinal binding pocket and W34 and W74 between two protomers are under 10 Å (Table S4). However, their chemical shifts deviate from the chemical shift observed for the observed cross peak. Thus, we exclude spin diffusion between nearby ^19^F atoms as origin of these cross peaks. Reinspection of the ^19^F spectrum on W34F (Fig. S6H) showed a small reduction in intensity at the two chemical shifts associated with the cross peak in the EXSY spectrum. Furthermore, the EXSY spectrum of [5-^19^F-Trp]-GPR-W34F is absent of cross-peaks (Fig. S14). We therefore conclude that these cross peaks arise from chemical exchange on the second timescale between two conformational states of W34.

To gain further insight into the two conformational states, two proteorhodopsin structures were compared (Fig. S15). The cryo-EM structure of GPR (pH 7.5)^38^ and the crystal structure of blue light-absorbing proteorhodopsin (BPR; PDB ID: 4KLY, pH 6.5)^29^ reveal distinct conformations of the highly conserved W34 residue. Notably, the amino acid sequences surrounding W34 are nearly identical in the two proteins, suggesting that the observed structural differences are not sequence-driven. To further investigate whether these conformations correspond to the chemical exchange observed experimentally, AF-QM/MM chemical shift predictions were performed using the BPR structure. The calculated ^19^F chemical shift for W34 in the BPR structure was -45.9 ppm (using the monomer with environmental charges), differing by 4.5 ppm from the value predicted for the cryo-EM GPR structure. This difference is of the same order of magnitude as the 2.7 ppm chemical shift difference observed in the EXSY spectrum. Together, these findings suggest that the exchanging states observed by NMR could be similar to the W34 conformations captured in the available structures.

Thereafter, we investigated the pH dependence of the exchange signals (Fig. S16). The cross-peak intensities decreased progressively with decreasing pH. This reduction may reflect a slowdown of the underlying exchange process or, alternatively, result from the overall signal broadening at low pH, which renders the exchange cross-peaks unobservable.

## DISCUSSION

### Biochemical characterization of [5-^19^F-Trp]-wtGPR

A small number of previous studies reported the use of 5-^19^F-Trp labelling of integral membrane proteins and their analysis by solution state NMR at fields of up to 600 MHz ^1^H Larmor frequency.^42–46^ The used membrane mimics ranged from organics solvents via detergent micelles to lipid nanodiscs. Here, we have utilized 5-^19^F-Trp labelling for proteorhodopsin which was then incorporated in liposomes. The demonstrated suitability of this labelling approach for MAS-NMR is in line with reports for soluble proteins.^12^ An additional benefit of testing this approach on a membrane bound photoreceptor is that the chromophore offers a highly sensitive experimental readout for the effect of 5-^19^F-Trp labelling in addition to standard biochemical analytics and NMR spectroscopy.

Metal affinity and SEC purification resulted in highly homogeneous samples and hNH experiments of [^15^N_2_-Lys]-labelled GPR show the same overall fold compared to nonfluorinated samples (Fig. 2C, S1C). [5-^19^F-Trp]-GPR also shows full proton pumping activity (Fig. 2B). The apparently larger pH gradient in whole cell assays compared with unlabeled GPR is likely a consequence of the lowered pK_a_ value of D97.

Nevertheless, some slight functional modifications of GPR have been observed. The UV-VIS spectra absorption maximum (λ_max_) exhibits an altered pH-dependence, reflecting a lower pK_a_ of D97 (Fig. 2A), which can also explain the increased pumping activity in the whole cell assays. This change in pKa is most likely due to the altered electrostatic environment caused by fluorination of W98, an integral component of the retinal binding pocket (Fig. S17) due to the inverted polarity of the C–F bond relative to the corresponding C–H bond in tryptophan.^47^ Furthermore, fluorination of W34 in the Trp34–His75–Asp97 triad may exert an allosteric effect on D97, potentially contributing to the observed changes. Equally intriguing is the reduced upper limit of the absorption maximum observed during the pH titration, which is also likely caused by changes in the electrostatic environment of the retinal chromophore induced by nearby 5-fluorotryptophan residues.

### Effect of MAS-rate and temperature

Ultrafast MAS at 100 kHz and above has been shown to efficiently disrupt the dipolar proton coupling network enabling direct proton detection.^9^ In case of sparse ^19^F labelling, homo- and heteronuclear dipole-dipole couplings are less dominant and MAS at 100 kHz should be sufficient to remove any residual couplings. Indeed, we observe a substantial improvement in line width with increasing MAS rate (Fig. 3A) which is in line with previous observations on solid, fluorinated pharmaceuticals^15^. Additional proton decoupling on these solid samples was shown to slightly improve resolution further.^48^ Here, deuteration of the most adjacent protons by using 5-^19^F-d_4_-Trp did not yield any further improvements indicating that sufficient decoupling through MAS in this membrane protein sample has been already achieved. The observed temperature dependence of the ^19^F line shapes also shows that molecular motions in GPR support efficient averaging.

### Chemical shift and lineshape analysis

Using tryptophan substitutions and substitutions of neighboring residues, assignment of all distinguishable signals was achieved due to sample rotation at 100 kHz (Fig. S6). In a previous study, ^19^F NMR at 60 kHz MAS allowed the distinction and assignment of five tryptophans in a viral capsid protein.^12^

Interestingly, W98 at -44 ppm falls outside of the chemical shift range of the other Trp signals. On this isolated signal, an overall global effect of all introduced mutations is seen (Fig. S6) illustrating the responsiveness of ^19^F towards even small or distant changes within the protein.

Spectral deconvolution revealed site-specific linewidth differences indicative of variations in side-chain mobility (Fig. 4A). Broader linewidths imply that the 5-^19^F-indole ring samples a wider conformational ensemble, experiences slower dynamics, or both. T_2_ measurements enabled the separation of homogeneous and heterogeneous contributions to the observed linewidth (Figs. 4B, S5). For all resonances, the homogeneous linewidth was markedly smaller than the total observed linewidth, indicating that heterogeneous broadening dominates the spectral linewidth for most sites, but there are interesting differences: W98 and W197 are both found within the retinal binding pocket (Fig. 1) with similar homogeneous linewidth, but W98 has a much larger heterogeneous contribution (Fig. 4B) indicating a higher degree of plasticity for this sidechain. The other two residues with large heterogeneous linewidth contributions are W149 and W239, which are exposed to the surface of GPR.

The known responsiveness of fluorine to its electronic environment led to a large observed chemical shift dispersion. This raised the question which determinants in GPR are responsible and how they can be estimated. To this end, we characterized the environment of each tryptophan residue using an empirical and intentionally simplified interaction score (Eq. 1) derived from the cryo-EM structure of GPR.^38^ Despite its simplicity, the resulting interaction score exhibits a reasonable linear correlation with the experimental ^19^F chemical shifts (Fig. 4).

For example, W98 exhibits both the highest interaction score and the most downfield ^19^F chemical shift, primarily owing to its proximity to the retinal Schiff base and Y95. The dominant contributions arise from charge–dipole interactions with the protonated Schiff base and dipole–dipole interactions with Y95, resulting in a dominating first term in our interaction score (Eq. 1). Consistent with this interpretation, substitution of Y95 with the nonpolar residue phenylalanine resulted in a pronounced upfield shift of the W98 resonance (Fig. 4C). Nevertheless, the ^19^F resonance of W98 remained the most downfield shift of all tryptophan residues in the Y95F variant, indicating that the retinal Schiff base is the dominant determinant of its chemical shift. Thus, the combined electrostatic and polar interactions within the local environment of W98 strongly modulate its ^19^F chemical shift.

In contrast, W149 and W239 exhibit the lowest interaction scores and among the most upfield chemical shifts. Both residues are located at the protein surface with their side chains oriented toward the lipid membrane, where they experience substantially fewer polar and electrostatic interactions than W98.

To assess whether and how accurately ^19^F chemical shifts can be predicted for a membrane protein of this size, we compared the experimental chemical shift dispersion with state-of-the-art quantum chemical calculations. Predicting ^19^F shifts in proteins has been pursued since the early 1990s using empirical shielding models combined with molecular dynamics.^49^ DFT-based Fluorine chemical shifts been computed for many years, but most studies focused on small fluorinated compounds.^50–53^ ^19^F chemical shifts of fluorinated tryptophans have been characterized in crystalline amino acid models by solid-state NMR and DFT,^40^ but predicting these shifts in the context of a large membrane protein requires methods that explicitly account for the surrounding protein and solvent environment. Cluster-DFT approaches with implicit solvation have been applied to smaller proteins,^54^ and QM/MM calculations have been used for ^19^F shifts of ligands bound to proteins,^41^ or labelled sites within proteins^54^ yet the prediction of chemical shifts for fluorinated amino acid residues within a membrane protein assembly remains challenging. To the best of our knowledge, no predications of 5-^19^F-Trp chemical shifts of different sites within a protein structure have been published. One major challenge is the large size of the system, which requires linear scaling approaches, which have been used, on case of ^19^F for predicting chemical shifts of protein-bound ligands^41^ or for ^13^C or ^15^N chemical shifts of membrane proteins^24, 30^. Here, the AF-QM/MM approach was employed. Remarkably, the calculated and experimental chemical shifts showed very good agreement and the chemical shift trend between the different sites is well reproduced (Fig. 4D). The remaining deviations likely arise from differences between the computational model and the experimental conditions, including pH, ionic strength, solvent treatment, and, importantly, the presence of a lipid environment. Recent work has shown that predicted electrostatic contributions to ^19^F chemical shifts of fluorotryptophan residues are substantially attenuated by dielectric shielding from the protein and solvent environment, complicating the quantitative prediction of site-specific shifts in globular proteins.^55^

Interestingly, only W83 emerged as a clear outlier. This residue is located near the water– membrane interface and may therefore be more sensitive than the other residues to differences between the computational model and the experimental environment. An alternative explanation is a local conformational difference between the cryo-EM structure used for the calculations and the conformational ensemble sampled under the NMR conditions. However, if this were the case, the interaction score A would also be expected to deviate significantly, which is not observed. The origin of this discrepancy therefore remains unclear and will require further investigation.

Despite this single outlier, the overall performance of the linear-scaling AF-QM/MM approach for site-resolved prediction of ^19^F chemical shifts in proteins is highly encouraging. The method has the potential to facilitate resonance assignment, improve the interpretation of ^19^F NMR spectra, and provide a valuable tool for discriminating between alternative structural models and experimental conditions.

### pH-dependent changes in GPR

Our ^19^F NMR spectra reveal a pronounced pH dependence. At low pH, all resonances undergo substantial line broadening (Fig. 5A). Previous studies have shown that acidic pH shifts GPR toward a more closed conformational state compared with higher pH values.^38^ Our data further demonstrate that this conformational transition is accompanied by a marked change in protein dynamics, as evidenced by the widespread line broadening. These findings suggest that acidification alters not only the average conformation of GPR but also its underlying conformational dynamics.

Due its distinct chemical shift and its high degree of conservation within the retinal binding pocket, W98 is discussed in detail. At pH 5, the W98 signal broadens dramatically. Its line shape is almost fully governed by inhomogeneous contributions in contrast to pH 9, which lets us conclude that the W98 side chain movement slows down from fast exchange to the intermediate timescale regime. The pronounced line broadening of the W98 resonance may be linked to the protonation state of the neighboring D97 residue (Fig. S16). Previous studies have shown that the photocycle slows at lower pH due to the protonation of D97.^26, 56^ This correlation raises the intriguing possibility that the highly conserved W98 residue acts as a molecular “pace-setter” for the conformational transitions underlying the photocycle. Specifically, fluctuations of W98 could provide a stochastic mechanism that governs transitions between conformational states in synchrony with photocycle progression.^57^ This pace-setting would require a coupling between W98 and the surrounding protein. Consistent with this hypothesis, the pH-dependent line broadening is not confined to W98 but is observed throughout the ^19^F spectrum of [5-^19^F-Trp]-GPR, indicating that the other tryptophan residues react similar to W98. Likewise, the assignment mutants reveal that substitutions at distant sites perturb the W98 resonance through changes in linewidth and/or chemical shift (Fig. S6), further supporting the existence of an allosterically coupled network involving W98. Although these observations are consistent with such a model, direct evidence for this coupling remains to be established. A promising approach would be to investigate GPR variants with altered D97 pK_a_ values, such as the H75N mutant, to determine whether perturbing D97 protonation correspondingly modulates the dynamics of W98.

At the protomer interface, W34 exhibited intriguing pH-dependent dynamics. Two-dimensional EXSY spectra revealed slow exchange of W34 at high pH (Figs. 5C, D), whereas no exchange was observed at low pH. Together with the available structural data and ^19^F chemical shift predictions, these findings suggest that W34 undergoes ring flipping on the seconds timescale at high pH. Based on the available structural information, previous studies of the Trp-His-Asp triad, and our observations, we propose the following hypothetical model (Fig. 6): At low pH, W34 is oriented toward H75 in the neighboring protomer. This interaction is proposed to stabilize H75 in a conformation in which it can form a hydrogen bond with D97^28^, thereby restricting the ring-flipping motion of H75.^27^ Consequently, proton transport is impaired under acidic conditions.^26, 58^ At higher pH, however, the W34–H75 interaction is transiently disrupted for several seconds, allowing increased conformational flexibility of H75 and thereby facilitating proton transport during these intervals. Because this disruption is only transient, proton transport is not continuously enhanced but rather appears to be “throttled.” Consistent with this model, previous ensemble transport measurements have shown reduced proton transport efficiency at higher pH.^26, 58^

**Figure 6:**
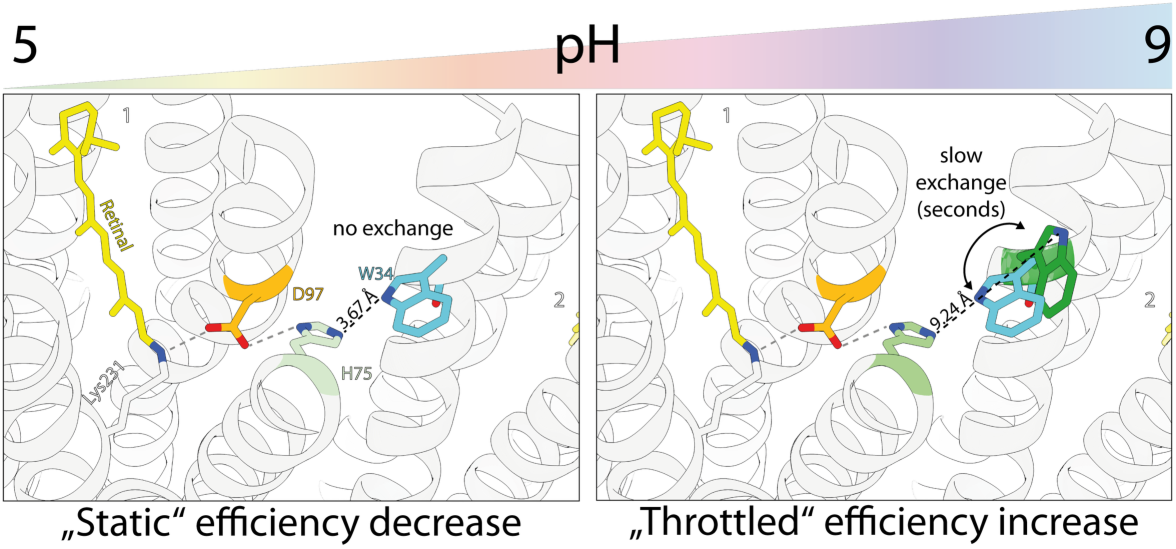
Model for interprotomeric pH-sensing by proteorhodopsins. At low pH, W34 performs no ring-flipping motion thus keeping contact to the neighboring H75. Subsequently proton transport is impaired under acidic conditions. At higher pH, a ring-flipping motion of W34 disrupts the contact to H75 in the seconds range thus throttling the proton transport in the neighboring protomer.

As this throttling appears to be modulated by pH, it may represent a pH-dependent regulatory mechanism in proteorhodopsin. Such a mechanism could prevent excessive alkalinization of the cell when the external environment becomes more alkaline by limiting proton transport under these conditions. In summary, the Trp-His-Asp triad may function as an interprotomeric, pH-dependent regulatory element that modulates proton transport through changes in protein dynamics rather than large-scale structural rearrangements. The observation of slow ring-flipping motions detected by ^19^F NMR was also reported in a recent work on flaviviral proteases using genetically encoded 5-fluorotryptophan^59^. More broadly, these findings illustrate that such temporal alterations in conformational dynamics, in addition to structural changes, can play a critical role in determining protein function.

## CONCLUSION

Our study demonstrates that ^19^F ultrafast MAS NMR provides novel insights into the functional mechanism of proteorhodopsin by resolving the dynamics of two key tryptophan residues, W34, a component of the interprotomer His-Asp-Trp triad, and W98 within the retinal-binding pocket. Through the combined analysis of line shapes, chemical shifts, and conformational exchange, we identify W34 as a dynamic regulator of proton transport and provide further evidence for its role in pH-dependent gating. We propose that the interprotomer His-Asp-Trp triad functions as a pH-dependent molecular throttle, in which slow ring flipping of W34 on the seconds timescale transiently modulates the interaction with H75 and thereby regulates proton transport. Such a mechanism may help prevent excessive alkalinization of the cell under alkaline environmental conditions. The multichannel readouts provided by GPR also demonstrated that the incorporation of 5-^19^F-Trp has minimal influence on the structure and function of this membrane protein, an aspect which needs to carefully considered in each case.

Beyond these mechanistic insights, this work establishes broadly applicable principles for investigating membrane protein dynamics by ^19^F ultrafast MAS NMR. We demonstrate the advantages of MAS rates of 100 kHz and beyond for resolving conformational heterogeneity and dynamics in complex membrane environments. Furthermore, the introduction of a simple scoring metric, together with the integration of AF-QM/MM calculations, facilitated the interpretation of ^19^F chemical shifts and enabled robust conformational assignments in a challenging membrane protein system.

More broadly, the workflow presented here combines ^19^F ultrafast MAS NMR with structure-based chemical shift calculations into a versatile framework for probing membrane protein structure and dynamics at atomic resolution. Given the exceptional sensitivity of ^19^F chemical shifts to the local molecular environment, this approach should be readily transferable to other rhodopsins with diverse functional properties as well as to pharmacologically important membrane proteins, including ABC transporters and GPCRs, particularly when investigated in native-like lipid assemblies.

## MATERIALS AND METHODS

### Mutagenesis

The GPR mutants W34F and W74F were generated previously.^27^ All other mutants were generated using Quikchange single-site directed mutagenesis using the primers in Table S5.

### Expression and purification

All constructs were expressed in *E*. *coli* C43(DE3) cells as described previously.^27^ Briefly, the cells were grown in defined medium at 37 °C. In order to incorporate 5-fluorotryptophan, defined medium lacking tryptophan was used for expression. When the culture reached an OD_600_ of 0.5, 50 mg/l culture of 5-fluoro-L-tryptophan or 100 mg/l culture of 5-fluoro-DL-tryptophan-2,4,6,7-[d4] were added to the medium and the culture was shaken for another 10 minutes until protein expression was induced with IPTG and retinal. After induction, the temperature was lowered to 27 °C and the cells were grown for another 16 h. After harvesting at 5000 × g for 10 min at 4 °C, the cell pellet was stored at -80 °C or directly lysed.

All GPR constructs were purified as described previously. For NMR spectroscopy, the eluates after immobilized metal affinity chromatography (IMAC) were additionally subdued to size-exclusion chromatography using a Superdex 200 increase 10/300. The peak containing the highest absorbance at 520 nm was isolated and used for reconstitution.

### Reconstitution

Reconstitution of all GPR constructs into liposomes containing 9:1 (w/w) 1,2-dimyristoyl-sn-glycero-3-phosphocholine and 1,2-dimyristoyl-sn-glycero-3-phosphatidic acid (DMPC/DMPA) was performed as described previously.^27^ In order to adjust the pH of the proteoliposomes, the latter was sedimented at 92,000 × g and 4 °C for 30 min. Then, the pellet was washed five times using either 50 mM Tris (pH 9), HEPES (pH 7.5), MES (pH 6) or acetate buffer (pH 5) using the same centrifugation conditions.

For NMR measurements, the proteoliposome suspension was further supplemented with 3 mM DSS and 0.5 mM TFA for calibration and then centrifuged into the MAS rotor using a home-built packing tool at 108,000 × g and 4 °C for 30 min.

### pH titration

After IMAC, the protein was rebuffered into pH titration buffer (50 mM Tris, 50 mM Na_2_HPO_4_, 50 mM boric acid, 50 mM sodium citrate, 100 mM NaCl, 0.05% DDM, pH 7.5) using a PD10 desalting column. After concentrating the protein to at least 1 mg/ml, either 4 M HCl or NaOH was titrated to the protein solution in approximately pH δ0.5 steps. The pH was measured and a small amount of protein was taken. After the titration, the absorbance spectra of the taken spectra were recorded using a ClarioStar plus (BMG Labtech) plate reader. The wavelength maxima of GPR at different pH were determined using the associated software package after curve smoothing using a moving average of seven.

### In-cell proton transport

After expression of GPR in C43(DE3) (see above), the cells were washed thrice using a salt solution consisting of (10 mM NaCl, 10 mM MgCl_2_, 1 mM CaCl_2_) by centrifugation at 2500 × g for 10 minutes. The transport assay was performed as described previously with the exception that transport was observed under three illumination cycles in which the cell suspension was illuminated for 4 min then rested for 3 min and illuminated again.

### NMR spectroscopy

All experiments were performed on a Bruker Avance III wide-bore ssNMR spectrometer operating at 800 MHz (^19^F Larmor frequency) equipped with a Bruker 0.7 mm HCN probehead, which was tuned on the proton channel to ^19^F. If not indicated otherwise, experiments were performed at 100 kHz MAS. In order to ensure a similar sample temperature for varying MAS frequencies, the proton chemical shift of water was monitored. Spectra were referenced to 0.5 mM buffered TFA or 3 mM DSS which was added before sedimentation into the MAS rotor.

hNH experiments were performed at approx. 25 °C. The temperature was calibrated by using the temperature-dependent ^1^H chemical shift of water.^60^ Chemical shift perturbations were calculated as the difference between the root mean squares of the chemical shifts.

^19^F spectra were acquired using a Hahn echo pulse sequence with an inter pulse delay of 100 µs in order to reduce background signals from the probehead. 90° pulse duration was 1.05 µs. Typically, 10,000 transients were acquired with an acquisition time of 50 ms. During processing, the FID was typically cut after 10 ms and 20 Hz exponential line broadening was applied.

^19^F EXSY were acquired with an acquisition time of 50 ms in the direct and 1.9 ms in the indirect dimension with 48 increments. 864 transients were recorded for each increment. Qsine (SSB = 4) and exponential (50 Hz line broadening) window functions were applied to the indirect and direct dimension, respectively.

Deconvolution of the processed 1D spectra was performed using OriginPro. Therefore, a Lorentzian was assumed for all peaks. For the T_2_ determination, the obtained linewidths from the initial deconvolution as well as chemical shifts were kept constant and applied to the residual spectra. The obtained peak heights and the corresponding errors from these deconvolutions were used and a monoexponential decay function was fitted through these points in order to obtain T_2_. Linewidths derived from T_2_ were calculated using the following equation:

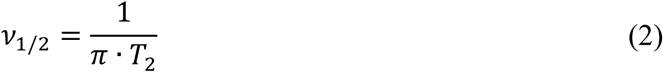

### 19F chemical shift calculation

In the AF-QM/MM approach employed this work, specific residues, 5-^19^F-Trp were designated as the core region. Residues within a predefined distance range from the core region constitute the buffer region, which is designed to capture the effects of the immediate local environment. Given that these local environmental effects are critical to the behavior of NMR chemical shifts, the buffer region was treated explicitly alongside the core region using QM (DFT) methods to ensure accurate modeling.

Computational models were constructed as follows: Cryo-EM structure of GPR (PDB ID: 7b03) served as the primary structural template for chemical shift predictions. The X-ray crystal structure of and BPR (PDB ID: 4kly) was additionally used to model the alternative conformation of W34. Two distinct models were designed: (1) monomer + solvent (one protomer extracted from the GPR pentamer structure); (2) pentamer + solvent. To further evaluate the long-range effects of environmental charges on the calculated fluorine chemical shifts, calculations were performed both with and without the environmental charges outside the buffer region.

The buffer region was defined using two distance-based criteria: (1) contacts between non-hydrogen atoms of the core and non-core residues within 5.0 Å; and (2) hydrogen-hydrogen interactions between the core and non-core regions within 2.5 Å. Boundary residues of the buffer region were subjected to hydrogen saturation to form complete closed-shell fragments. Both the core and buffer regions were treated with QM methods, while the remainder of the system was represented by background charges to model the electrostatic environment. Atomic point charges were derived from AMBER force field parameters. Further details of AF-QM/MM approach can be found in our previous work.^30^

^19^F chemical shifts were calculated using the density functional theory (DFT) gauge-including atomic orbital (GIAO) method implemented in the Gaussian 16 package.^61^ All fragment calculations were performed in parallel on a Linux cluster equipped with 24-core Intel Xeon 2.60 GHz processors. Isotropic chemical shifts were defined as the difference between the isotropic shielding of the reference (TFA) and the calculated isotropic shielding of the target system. This ensures that both experimental and calculated ^19^F chemical shifts are reported on the same TFA reference scale. The density functional and basis set employed for NMR calculations were B3LYP and 6-311G**, respectively, with the GD3BJ empirical dispersion correction applied.

## SUPPORTING DATA

Supporting biochemical data (Fig. S1) and topology plot (Fig. S2), additional NMR data (Figs. S3 and S4), supporting data explaining resonance assignment (Figs. S5, S6, S7), additional computational data (Figs. S8, S9 and S10), supporting pH-dependent NMR data (Figs. S11, S12, S13), comparison of PR structures at different pH values (Fig. S14), pH-dependent EXSY spectra (Fig. S15), illustration of polar environment of W98 (Fig. S16), spectra deconvolution results (Tables S1, S2), statistical analysis of chemical shift predictions (Table S3), distances between ^19^F atoms (Table S4), list of primers (Table S5).

## ACKNOWLEDGEMENTS

This work was funded by the LOEWE project GLUE (G protein-coupled receptor Ligands for Underexplored Epitopes) of the state of Hessen and by the CRC 1507 ‘Membrane-associated protein assemblies, machineries and supercomplexes’. D.D. was supported by the National Natural Science Foundation of China (Grant No. 22403029) and the China Postdoctoral Science Foundation (Grant No. 2024M760922). X.H. was supported by the Shanghai Municipal Science and Technology Commission with Grant No. 25511102400, National Natural Science Foundation of China (Grant Nos. 92477103 and 22273023), the Shanghai Frontiers Science Center of Molecule Intelligent Syntheses, and the Fundamental Research Funds for the Central Universities. We also acknowledge the Supercomputer Center of East China Normal University (ECNU Multifunctional Platform for Innovation 001) for providing computer resources. We thank Dr. Jagdeep Kaur, Frankfurt for helpful discussions.

## Notes

### Competing Interest Statement

The authors have declared no competing interest.

