## Supporting Information for "^19^F Ultrafast MAS NMR Reveals the Dynamic Basis of pH-Dependent Regulation in Proteorhodopsin"

### 1. Supporting biochemical data

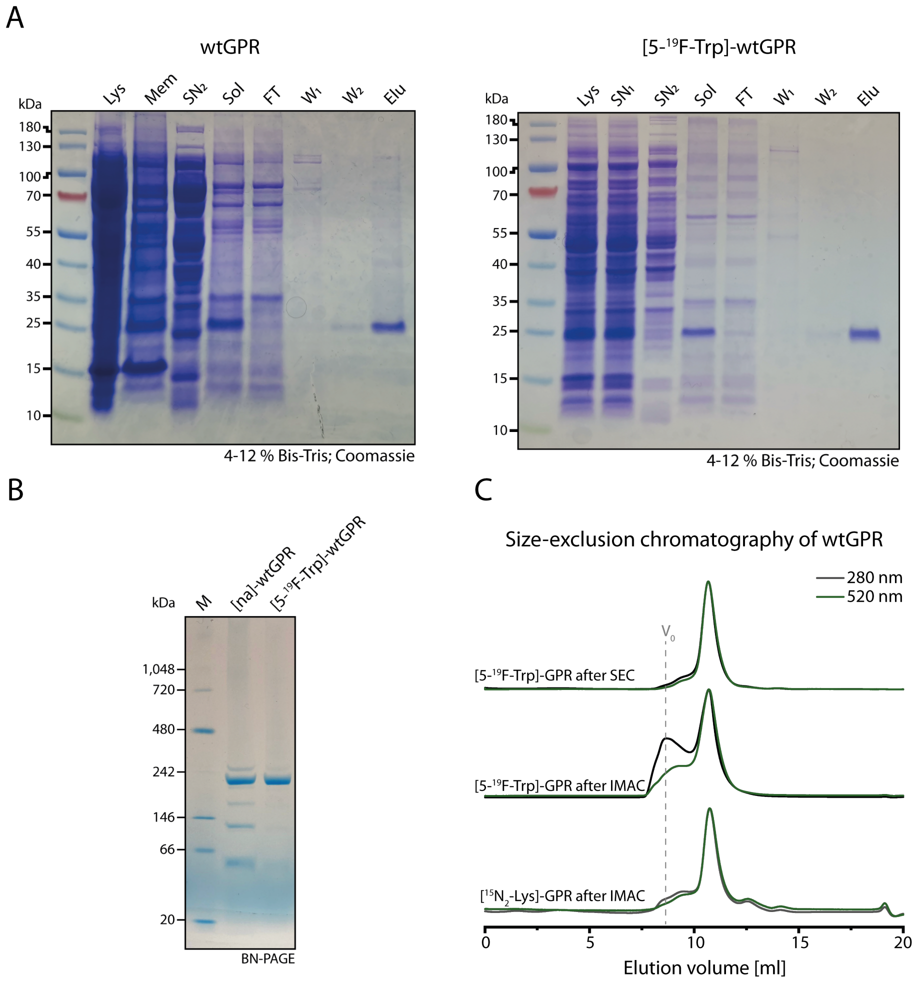


**Figure S1:** Biochemical characterization of [5-$^{19}$F-Trp]-wtGPR. **(A)** SDS-PAGE of IMAC purification of unlabeled [5-$^{19}$F-Trp]-wtGPR. Samples of different purification steps were applied on the SDS gel. Lys: after cell disruption; $SN_1$: after first centrifugation; $SN_2$: supernatant after ultracentrifugation; Mem: resuspended membranes; Sol: after solubilization; FT: flow-through; $W_1$: first wash fraction; $W_2$: second wash fraction; Elu: eluate. **(B)** BlueNative-PAGE of unlabeled and [5-$^{19}$F-Trp]-wtGPR. **(C)** Size exclusion chromatography of [5-$^{19}$F-Trp]-wtGPR after different purification steps (middle and top traces) compared to [$^{15}N_2$-Lys]-wtGPR (bottom trace). During SEC, absorption at 280 (black) nm and 520 nm of the bound retinal (green) was monitored. The void volume $V_0$ is indicated with a grey dashed line.

After IMAC purification, both [5-$^{19}$F-Trp]-wtGPR and [$^{15}$N-Lys]-wtGPR samples were analyzed by size-exclusion chromatography (SEC). Both preparations exhibited a narrow peak at the same elution volume, with absorbance at 280 nm arising from Trp or 5-$^{19}$F-Trp residues and at 520 nm from the retinal chromophore (Fig. S1C, middle and bottom). In addition, a peak was observed in the void volume ($V_0$). This peak was substantially more pronounced for the [5-$^{19}$F-Trp]-wtGPR sample and showed stronger absorbance at 280 nm than at 520 nm, indicating the presence of protein aggregates that lacked bound retinal (Fig. S1C, middle). The corresponding fraction was collected and exhibited visible precipitation after one day.

In contrast, non-fluorinated GPR preparations such as [$^{15}$N-Lys]-wtGPR displayed only minor aggregate formation, remained free of visible precipitation for several days, and could be used directly for NMR experiments without additional SEC purification. For [5-$^{19}$F-Trp]-wtGPR, an additional SEC purification step was therefore performed by collecting the monomeric fraction eluting at 11 mL. After storage for one day, this fraction was reapplied to the SEC column and yielded a single monodisperse peak with the expected absorbance at 520 nm and no detectable aggregate peak (Fig. S1C, top). Consequently, all NMR experiments were performed using [5-$^{19}$F-Trp]-wtGPR samples subjected to this additional SEC purification step, resulting in highly homogeneous protein preparations.

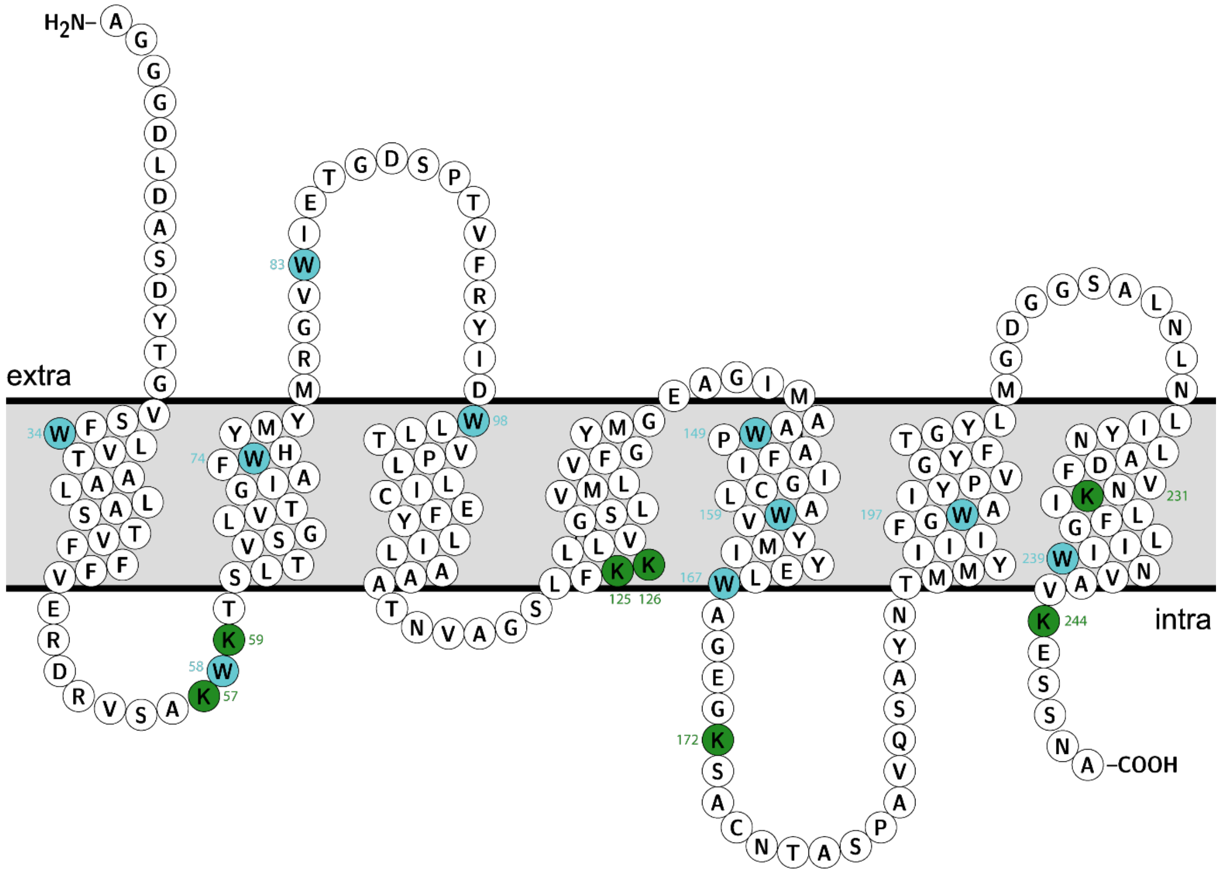


**Figure S2:** Membrane topology of GPR. Lysine residues are highlighted in green and tryptophan residues in blue.

#### 2. Supporting NMR data

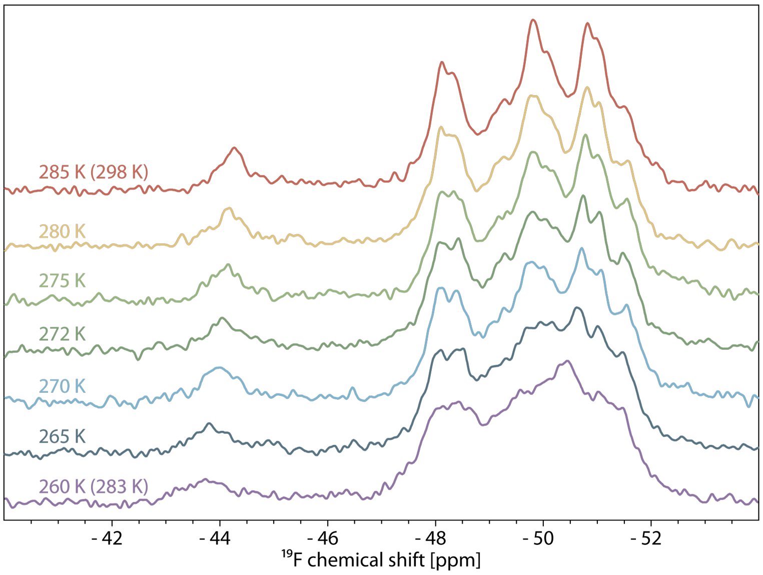


**Figure S3:** Temperature scan of [5-$^{19}$F-Trp]-wtGPR. $^{19}$F spectra were recorded at increasing nominal temperatures (bottom to top). The actual sample temperature was estimated *via* $^{1}$H chemical shift of water. The highest and lowest temperature correspond to 298 K and 283 K actual sample temperature, respectively.

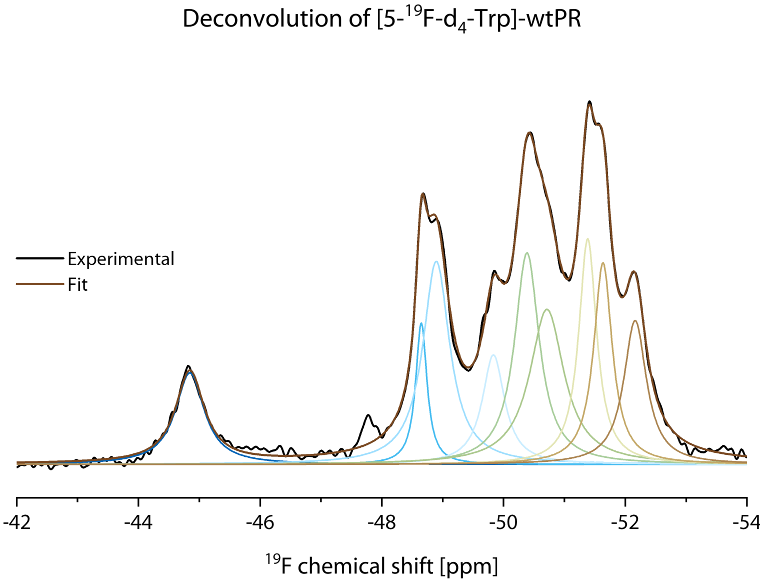


**Figure S4:** Spectral deconvolution of 1D $^{19}$F Hahn echo spectrum of [5-$^{19}$F-d$_4$-Trp]-wtGPR at pH 9 in DMPC/DMPA liposomes recorded at 298 K and 100 kHz MAS. Comparably to [5-$^{19}$F-Trp]-wtGPR, nine Lorentzian peaks were used to deconvolute the spectrum.

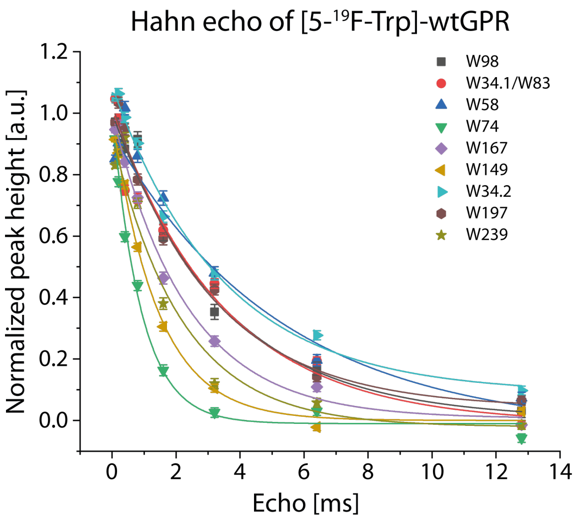


**Figure S5:** $T_2$ determination of [5-$^{19}$F-Trp]-wtGPR. Hahn echo with variable delays were recorded. The acquired spectra were deconvoluted with fixed chemical shifts and linewidth derived from the initial deconvolution (Fig. 4A).

### 3. Assignment

The assignment of all $^{19}$F resonances of [5-$^{19}$F-Trp]-wtGPR was established using site-directed mutagenesis (Fig. S6).

- **W98:** W98 was assigned based on the proximity mutation Y95F, which induced a chemical shift perturbation of the corresponding resonance (Fig. 4C). A direct W98F mutant could not be analyzed because the protein was unstable.
- **W167:** The $^{19}$F spectrum of W167F lacks the resonance at −49.83 ppm (Fig. S6A), which was therefore assigned to W167.
- **W58:** In the W58F spectrum, the resonance at −48.35 ppm is absent (Fig. S6B), allowing its assignment to W58.
- **W74:** The resonance at −49.23 ppm is missing in the W74F spectrum (Fig. S6C) and was therefore assigned to W74.
- **W197:** W197 was assigned based on the chemical shift perturbation of the resonance from −51.00 to −51.33 ppm in the I193T mutant (Fig. S6D).
- **W239:** In the W239F mutant, the most upfield-shifted resonance at −51.51 ppm is absent (Fig. S6E), allowing its assignment to W239.
- **W149:** The W149F spectrum shows a reduction in signal intensity between −49.00 and −50.25 ppm (Fig. S6F). Since the resonance of W167 had already been assigned, the remaining resonance in this region was assigned to W149.
- **W83:** The W83F mutant exhibits reduced intensity of the resonance at −48.12 ppm (Fig. S6G), which was therefore assigned to W83.
- **W34:** The W34F spectrum displays a decrease in signal intensity between −50.50 and −51.30 ppm. As the resonance at −51.00 ppm had already been confidently assigned to W197, the remaining resonance in this region was assigned to W34. In addition, the resonance at −48.12 ppm disappears completely (Fig. S6H). Based on the $^{19}$F–$^{19}$F EXSY spectra of wtGPR (Fig. 5C) and W34F (Fig. S14), this resonance was also assigned to W34, indicating that W34 gives rise to two exchanging resonances.
- **W159:** The W159F spectrum exhibits a general loss of spectral resolution (Fig. S7). The W159 resonance may therefore be broadened beyond detection. Alternatively, it may contribute to the unresolved signals between −47 and −52 ppm, precluding an unambiguous assignment.
- **Unspecific signal P:** An additional resonance at −47.76 ppm was observed in all spectra with variable intensity. Control measurements of unlabeled GPR and the protease inhibitor cocktail indicated that this signal originates from nonspecific binding of the fluorinated protease inhibitor AEBSF to the protein (Fig. S8).

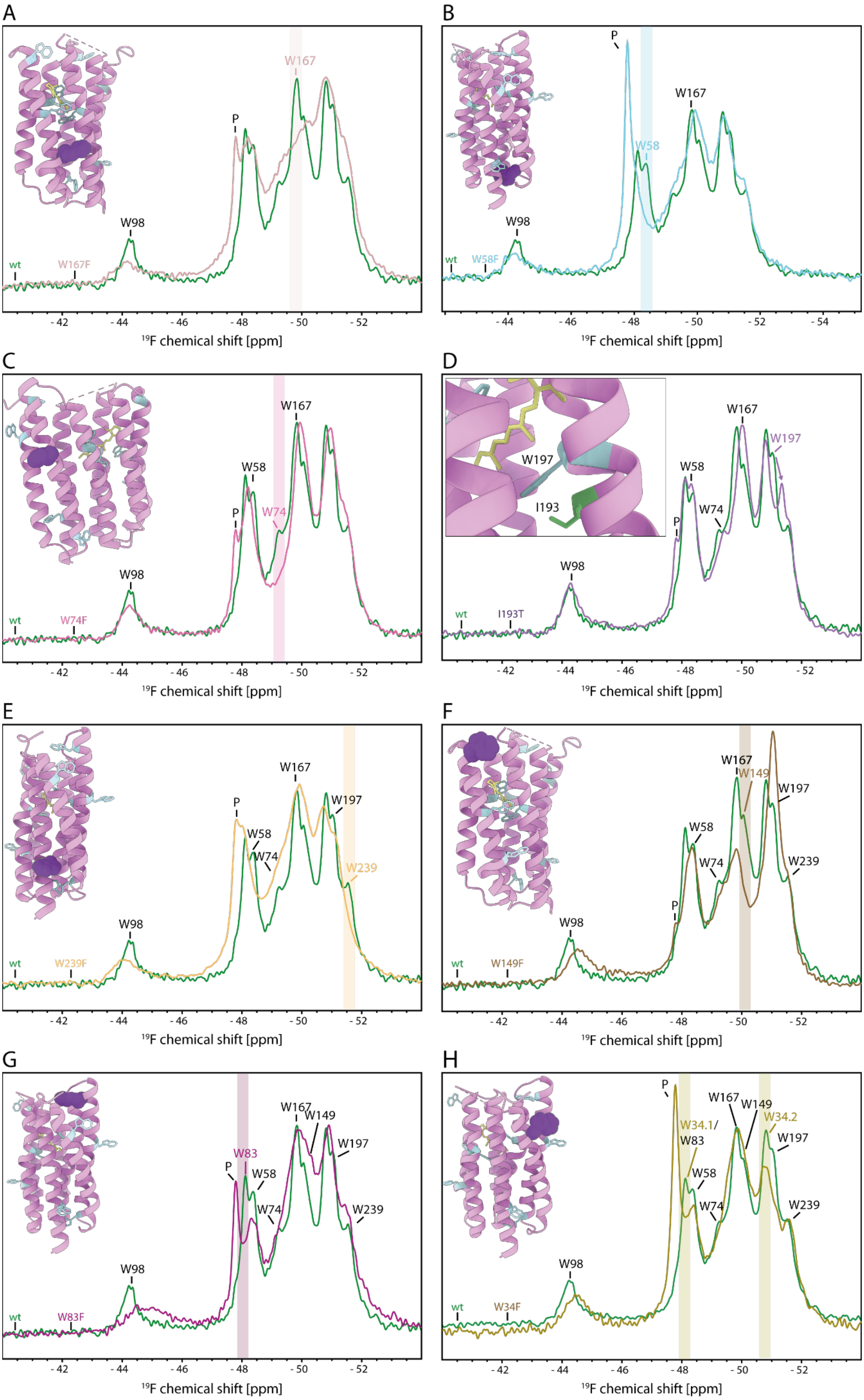

A
W167
P
W98
wt
W167F
- 42
- 44
- 46
- 48
- 50
- 52
¹⁹F chemical shift [ppm]
B
P
W167
W58
W98
wt
W58F
- 44
- 46
- 48
- 50
- 52
- 54
¹⁹F chemical shift [ppm]
C
W167
W58
P
W74
W98
wt
W74F
- 42
- 44
- 46
- 48
- 50
- 52
¹⁹F chemical shift [ppm]
D
W197
I193
W167
W197
W58
W74
P
W98
wt
I193T
- 42
- 44
- 46
- 48
- 50
- 52
¹⁹F chemical shift [ppm]
E
W167
P
W197
W58
W74
W239
W98
wt
W239F
- 42
- 44
- 46
- 48
- 50
- 52
¹⁹F chemical shift [ppm]
F
W167
W149
W197
W58
W74
W239
P
W98
wt
W149F
- 42
- 44
- 46
- 48
- 50
- 52
¹⁹F chemical shift [ppm]
G
W167
W149
W197
W83
P
W58
W74
W239
W98
wt
W83F
- 42
- 44
- 46
- 48
- 50
- 52
¹⁹F chemical shift [ppm]
H
P
W167
W149
W34.1/
W83
W34.2
W197
W58
W74
W239
W98
wt
W34F
- 42
- 44
- 46
- 48
- 50
- 52
¹⁹F chemical shift [ppm]

**Figure S6:** Assignment of [5-$^{19}$F-Trp]-wtGPR based on site-directed mutagenesis. $^{19}$F spectra of W167F **(A)**, W58F **(B)**, W74F **(C)**, I193T **(D)**, W293F **(E)**, W149F **(F)**, W83F **(G)**, W34F **(H)** are shown in overlap with the spectrum of [5-$^{19}$F-Trp]-wtGPR. The spheres in the inset structure cartoon display the positions of the mutated tryptophans except for **(D)** where the local environment is highlighted. Colored bars indicate a missing or shifting signal in the mutant compared to the wildtype spectrum. The subsequently assigned resonances are then highlighted in red. Black colored residues correspond to the assignment form the preceding spectrum. "P" indicates the signal arising from residual protease inhibitor. The assignment is further explained in the supporting text above.

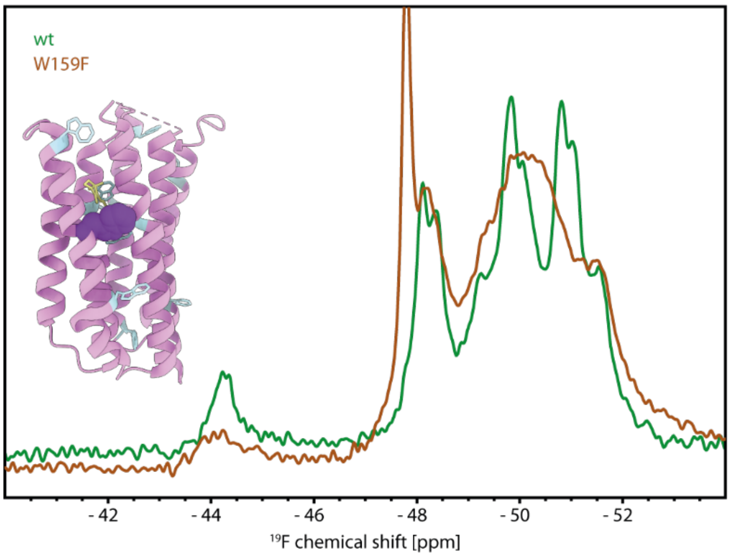


**Figure S7:** $^{19}$F spectrum of [5-$^{19}$F-Trp]-GPR-W159F (brown) overlayed with the wtGPR spectrum (green). The spheres in the inset structure displays the position of the mutated tryptophan.

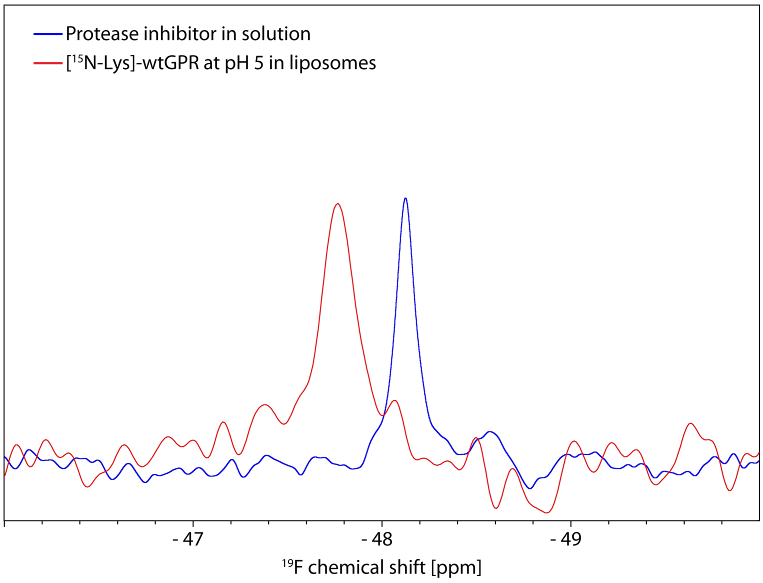


**Figure S8:** 1D $^{19}$F solution NMR spectra of the used protease inhibitor cocktail (blue spectrum) and $^{19}$F Hahn echo of [$^{15}N_2$-Lys]-wtGPR in liposomes at 100 kHz MAS (red spectrum). The signal stems from 4-(2-aminoethyl)benzenesulfonyl fluoride hydrochloride (AEBSF).

### 4. Supporting computational data

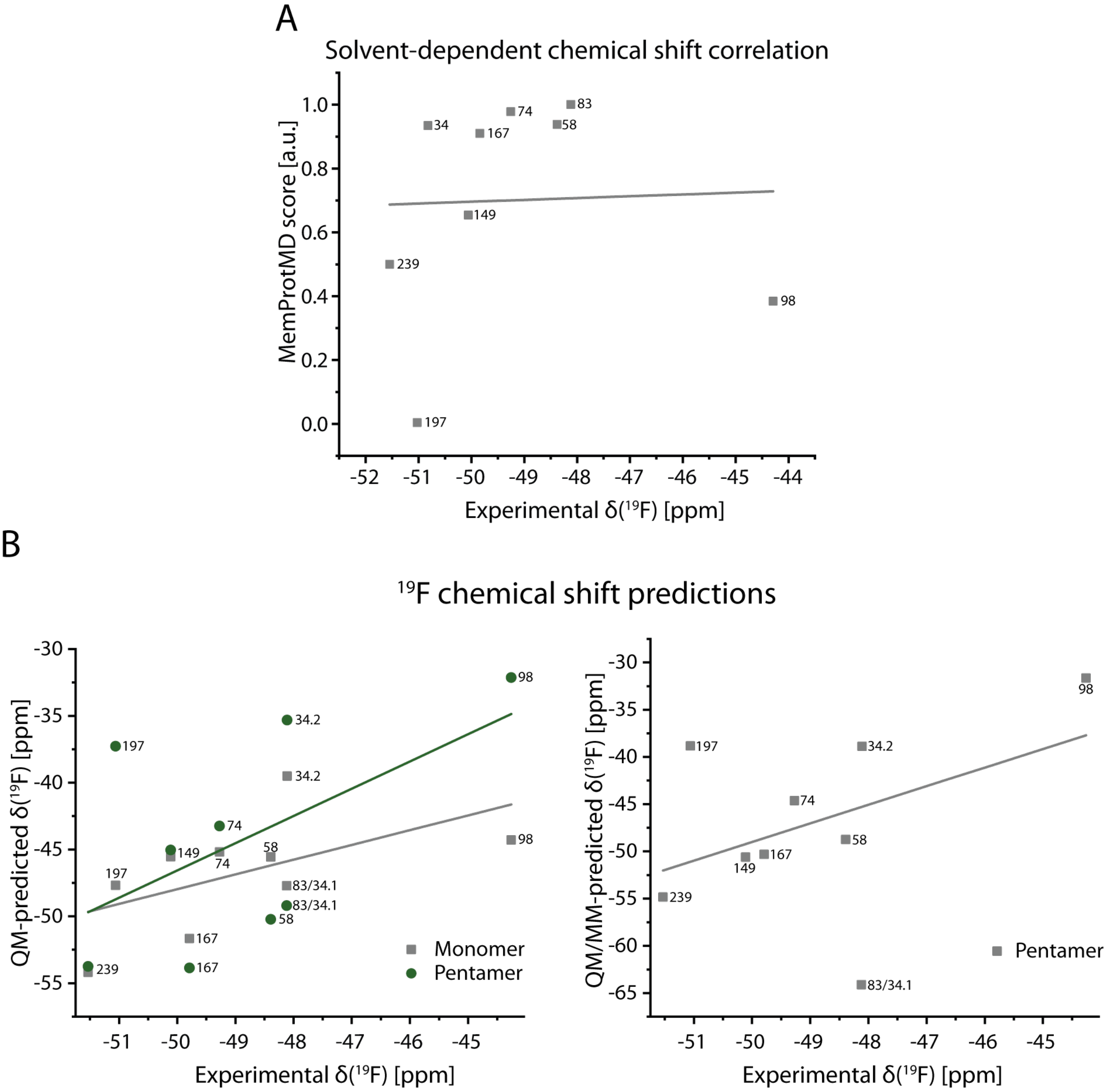


**Figure S9:** Estimating the chemical shift dispersion in $^{19}$F NMR. **(A)** Solvent scores from the MemProtMD database for tryptophan residues in the GPR cryo-EM structure (PDB ID: 7b03) are correlated with the experimentally determined chemical shifts of [5-$^{19}$F-Trp]-wtGPR. Numbers indicate the tryptophan residue. **(B)** Computational chemical shift predictions of [5-$^{19}$F-Trp]-wtGPR based on the cryo-EM structure (PDB ID 7b03). Left graph shows the predicted chemical shifts considering only the QM region. Chemical shift was predicted either considering only one monomer or the whole pentameric structure. Right graph shows the AF-QM/MM calculated chemical shifts of [5-$^{19}$F-Trp] with environmental charges included for the pentameric structure. The predictions for the monomer calculations are shown in Figure 4D.

Because no clear guidelines exist for selecting the buffer-region cutoff in fluorinated protein systems, we systematically tested cutoff distances from 3 to 6 Å on a GPR monomer. Both the Pearson correlation coefficient and RMSE indicated that the predicted $^{19}$F chemical shifts exhibited minimal variation when the cutoff distance was ≥ 4 Å (Fig. S9), so a 5 Å cutoff was chosen to balance accuracy and computational cost. We also evaluated the inclusion of ions in the model at the 3 Å cutoff, but this did not improve prediction accuracy (Fig. S10), so ions were excluded from all subsequent calculations.

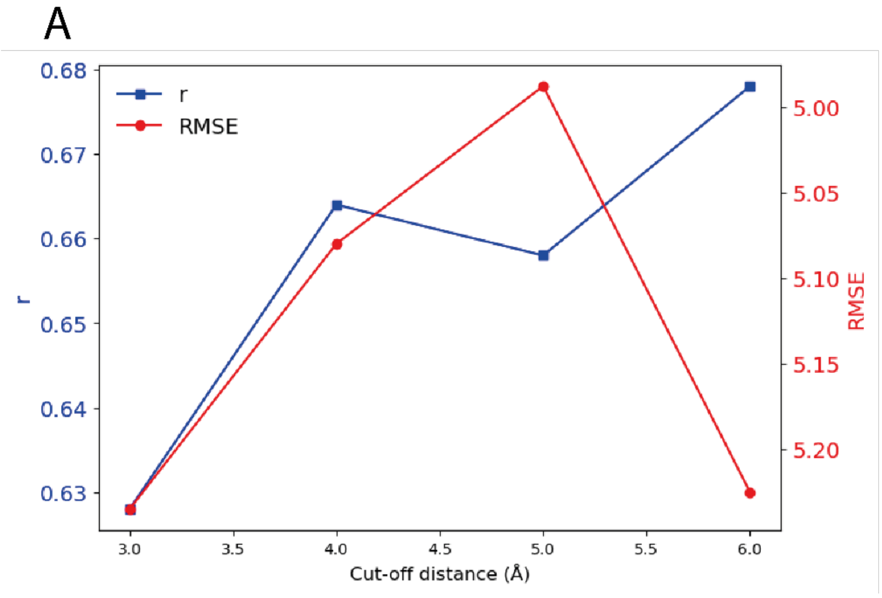


B

| Distance (Å) | r | RMSE |
|---|---|---|
| 3 | 0.628 | 5.235 |
| 4 | 0.664 | 5.080 |
| 5 | 0.658 | 4.988 |
| 6 | 0.678 | 5.225 |

**Figure S10:** Comparison of the Pearson correlation (r) and RMSE with different cut-off distance for buffer region in AF-QM/MM approach. **(A)** Overlay of r and RMSE in dependence of the Cut-off distance. **(B)** Values depicted in (A).

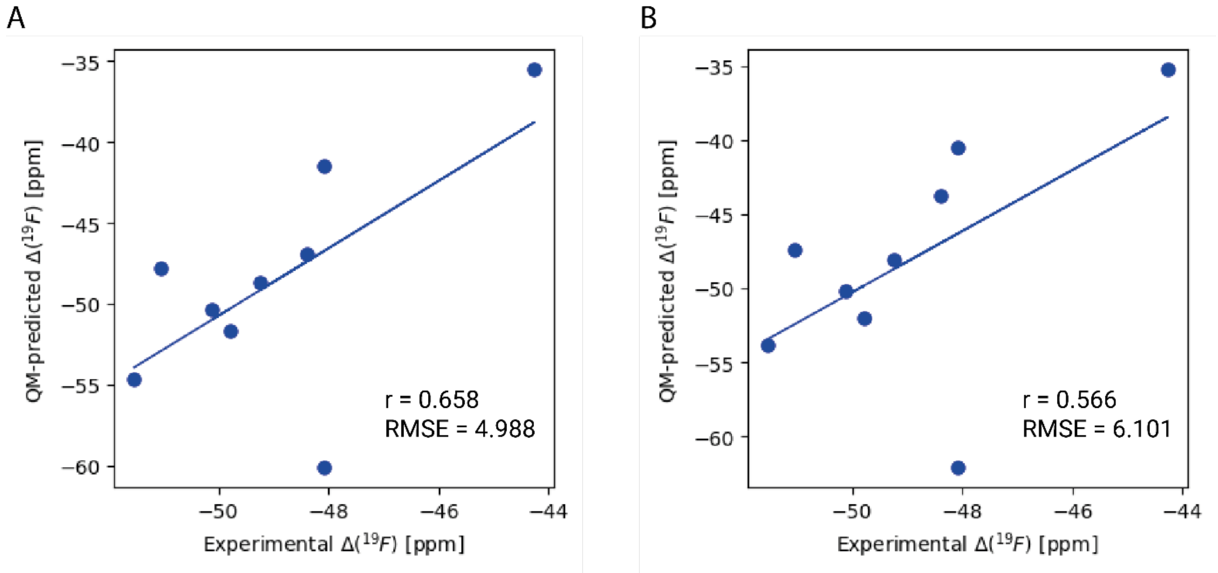


**Figure S11:** Comparison of Pearson correlation coefficient (r) and RMSE for $^{19}F$ chemical shift predictions in GPR monomer + solvent systems with a 3 Å cutoff distance, considering the presence **(A)** or absence of ions **(B)**.

### 5. Supporting pH-dependent data

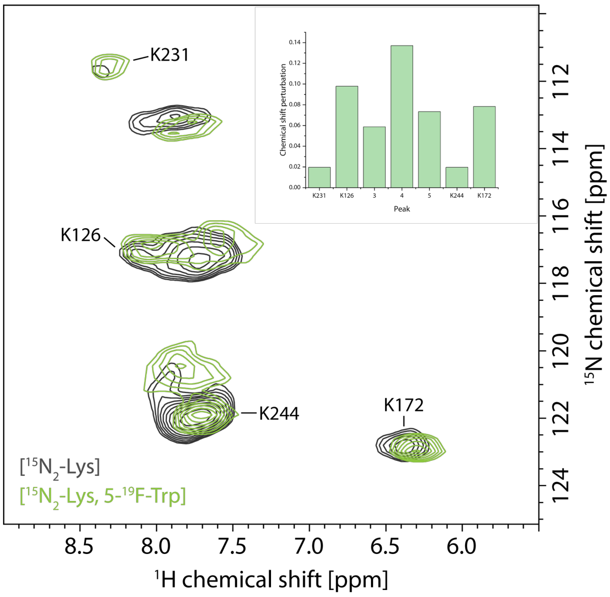


**Figure S12:** Proton-detected fingerprint spectra of labelled GPR at pH 5. hNH at 100 kHz MAS of [$^{15}N_2$-Lys]-wtGPR (black) and [5-$^{19}$F-Trp]-wtGPR (green) reconstituted in liposomes. Lysine residue assignments from previous assignments[1] are indicated. The insert shows chemical shift perturbations of the lysines between the two preparations.

A

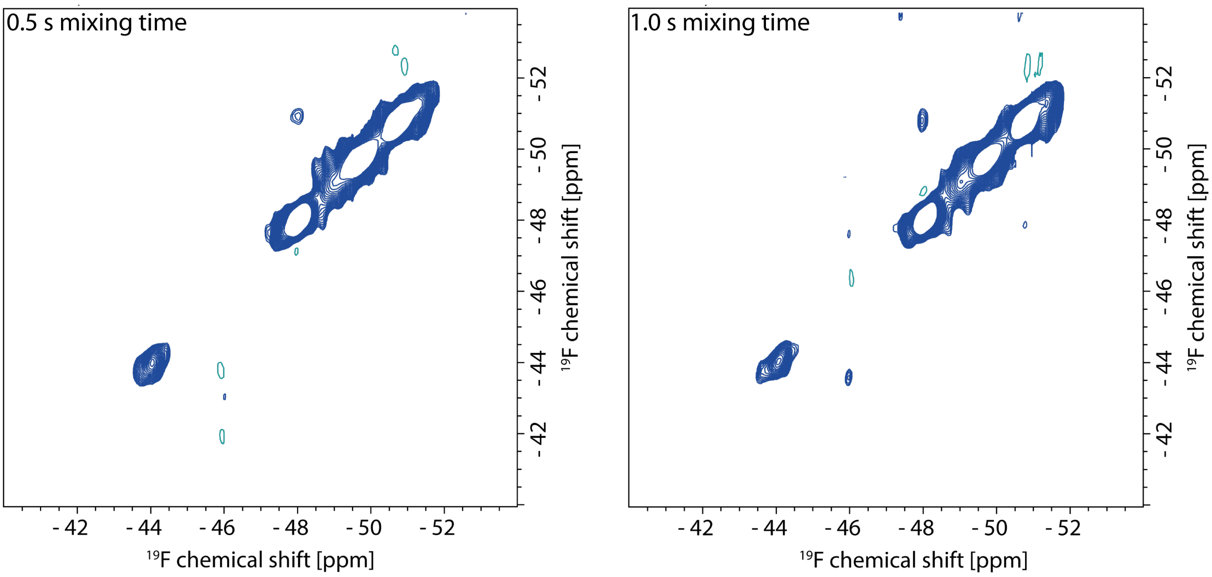


B

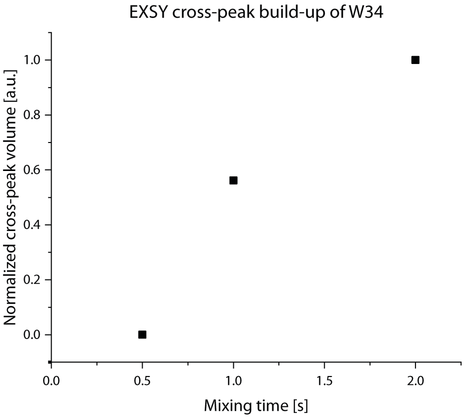


**Figure S13:** Supporting $^{19}F$-$^{19}F$ EXSY spectra. (**A**) $^{19}F$-$^{19}F$ EXSY of [5-$^{19}F$-Trp]-wtGPR at 0.5 and 1.0 s mixing times (see also Fig. 5C). **(B)** Normalized cross-peak volumes as a function of EXSY mixing times.

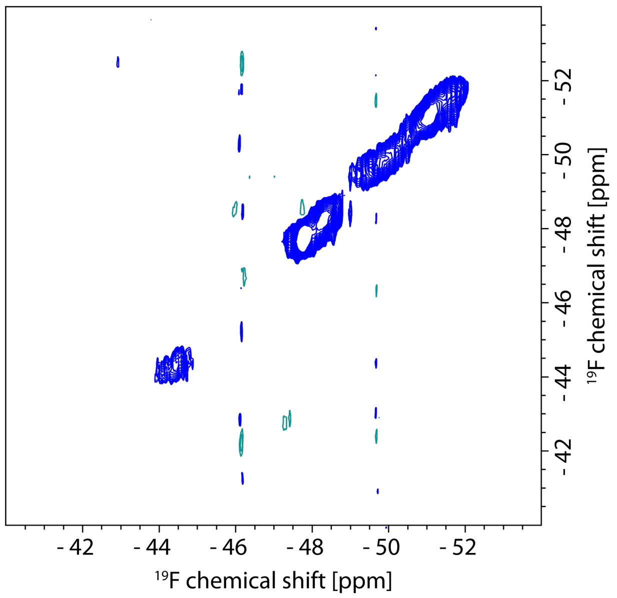


**Figure S14:** $^{19}$F-$^{19}$F EXSY spectrum of [5-$^{19}$F-Trp]-GPR-W34F with 2 s mixing time. No cross-peaks were observed.

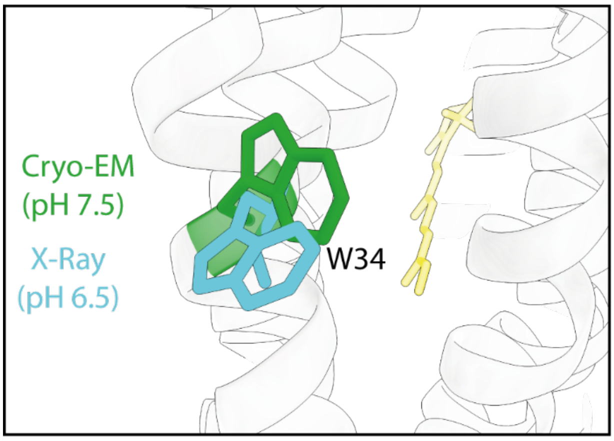


**Figure S15:** Conformations of W34. Comparison of the W34 from the cryo-EM structure (PDB 7b03; green)[2] with the crystal structure of BPR (PDB 4kly; blue)[3] by overlaying mentioned structures. The structures were prepared under different pH conditions.

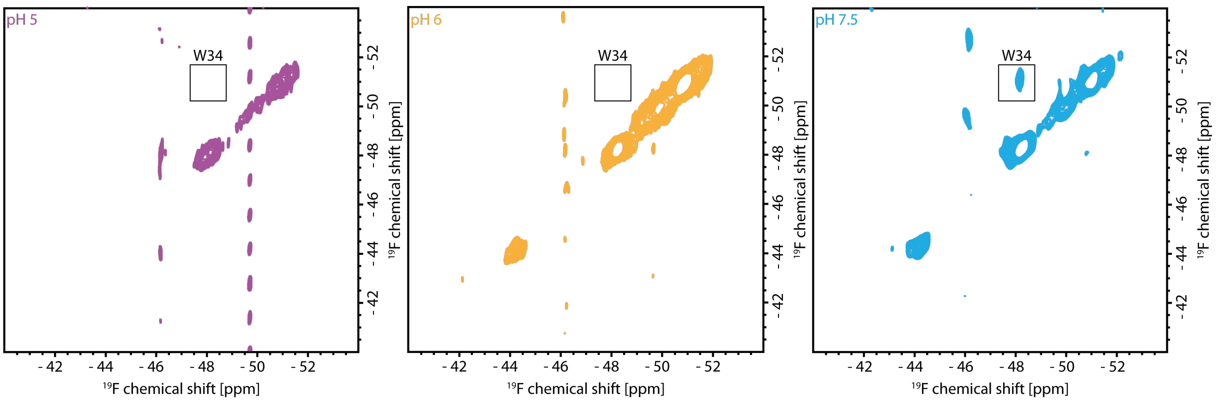


**Figure S16:** pH-dependent $^{19}F$-$^{19}F$ EXSY spectra. EXSY spectra were recorded at different pH values at 100 kHz MAS. The mixing time was 2 s. The temperature was approximately 25 °C.

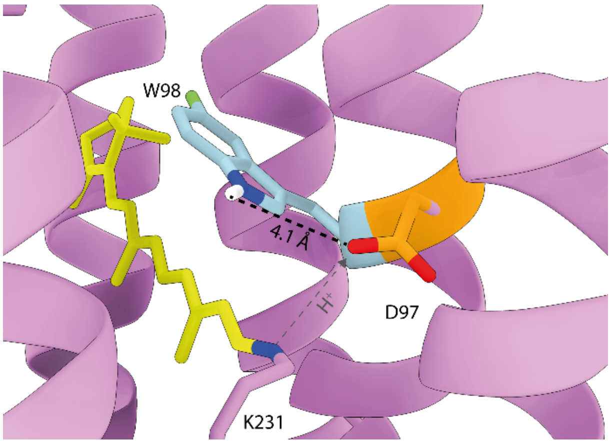


**Figure S17:** The polar environment of W98. W98 is an integral part of the retinal pocket. During the photocycle, the adjacent D97 accepts a proton from the Schiff base. Lowering the sample pH emulates this state.

### 6. Tables

**Table S1:** Homogeneous (**$\nu_{1/2,T2}$**) and observed full linewidth (**$\nu_{1/2,observed}$**) of [5-$^{19}$F-$d_4$-Trp]-wtGPR as well as chemical shifts and intensities derived from the deconvolutions and $T_2$ measurements.

| Peak | δ $^{19}$F [ppm] | $\nu_{1/2,observed}$ [Hz] | Observed intensity | $\nu_{1/2,T2}$ [Hz] |
|---|---|---|---|---|
| **W98** | -44.85 | 459 ± 8 | 282854 ± 3085 | 98 ± 13 |
| **W34.1/W83** | -48.65 | 180 ± 7 | 437922 ± 12877 | 106 ± 10 |
| **W58** | -48.89 | 428 ± 7 | 627146 ± 9546 | 53 ± 6 |
| **W74** | -49.83 | 357 ± 10 | 338359 ± 5080 | 92 ± 9 |
| **W167** | -50.38 | 392 ± 16 | 653239 ± 38471 | 256 ± 31 |
| **W149** | -50.71 | 547 ± 30 | 478911 ± 31537 | 166 ± 16 |
| **W34.2** | -51.38 | 285 ± 9 | 696679 ± 21092 | 265 ± 29 |
| **W197** | -51.63 | 294 ± 10 | 622534 ± 18835 | 69 ± 6 |
| **W239** | -52.16 | 361 ± 6 | 444927 ± 4390 | 139 ± 41 |

**Table S2:** Homogeneous (**$\nu_{1/2,T2}$**) and observed full linewidth (**$\nu_{1/2,observed}$**) of [5-$^{19}$F-Trp]-wtGPR as well as chemical shifts and intensities derived from the deconvolutions and $T_2$ measurements.

| Peak | δ $^{19}$F [ppm] | $\nu_{1/2,observed}$ [Hz] | Observed intensity | $\nu_{1/2,T2}$ [Hz] |
|---|---|---|---|---|
| **W98** | -44.26 | 396 ± 8 | 165612 ± 2166 | 90 ± 19 |
| **W34.1/W83** | -48.09 | 256 ± 4 | 423998 ± 4991 | 88 ± 24 |
| **W58** | -48.39 | 278 ± 7 | 357264 ± 4497 | 70 ± 16 |
| **W74** | -49.24 | 544 ± 14 | 240874 ± 2873 | 343 ± 38 |
| **W167** | -49.79 | 283 ± 9 | 452998 ± 14725 | 144 ± 13 |
| **W149** | -50.11 | 533 ± 18 | 395830 ± 9686 | 231 ± 17 |
| **W34.2** | -50.79 | 260 ± 6 | 512231 ± 7786 | 99 ± 10 |
| **W197** | -51.06 | 254 ± 8 | 382994 ± 7551 | 102 ± 16 |
| **W239** | -51.53 | 495 ± 8 | 307910 ± 2882 | 146 ± 40 |

**Table S3:** Pearson correlation coefficients for the linear regressions performed during the chemical shift predictions. The highest values are highlighted. The value in brackets corresponds to the correlation coefficient when W83 is not considered for the regression.

| Prediction | Correlation coefficient |
|---|---|
| Solvent score of MemProtMD | 0.03607 |
| Structure-based interaction score A | **0.77751** |
| AF-QM/MM (Monomer, buffer region) | 0.55788 |
| AF-QM/MM (Pentamer, buffer region) | 0.54383 |
| AF-QM/MM (Pentamer, whole protein) | 0.71942 |
| AF-QM/MM (Monomer, whole protein) | **0.63022 (0.91544)** |

**Table S4:** Distances between fluorine atoms in [5-$^{19}$F-Trp]-wtGPR. for distance determination, the fluorine atoms were modelled into the cryo-EM structure.

| Distance pair | Distance [Å] |
|---|---|
| W34 – W74 | 7.9 |
| W159 – W197 | 9.9 |

**Table S5:** List of primers used for Quikchange mutagenesis.

| Mutation | Primer |
|---|---|
| W58F | 5'tgaaagagatagagtttctgcaaaattcaaaacatcattaactgtatctggtc-3'<br>5'-gaccagatacagttaatgatgttttgaattttgcagaaactctatctctttca-3' |
| W83F | 5'-catgtacatgagaggggtattcattgaaactggtgattcgc-3'<br>5'-gcgaatcaccagtttcaatgaatacccctctcatgtacatg-3' |
| Y95F | 5'-tgattcgccaactgtatttagattcattgattggttactaacagttc-3'<br>5'-gaactgttagtaaccaatcaatgaatctaaatacagttggcgaatca-3' |
| W149F | 5'-gaagcaggaatcatggctgcattccctgcattcattattg-3'<br>5'-caataatgaatgcagggaatgcagccatgattcctgcttc-3' |
| W159F | 5'-gcattcattattgggtgtttagctttcgtatacatgatttatgaattattcg-3'<br>5'-cgaataattcataaatcatgtatacgaaagctaaacacccaataatgaatgc-3' |
| W167F | 5'-gtttagcttgggtatacatgatttatgaattattcgctggagaaggaaaatc-3'<br>5'-gattttccttctccagcgaataattcataaatcatgtatacccaagctaaac-3' |
| I193T | 5'-cagcttacaacacaatgatgtatattaccatctttggttgggc-3'<br>5'-gcccaaccaaagatggtaatatacatcattgtgttgtaagctg-3' |
| W239F | 5'-ctgactttgttaacaagattctatttggtttaattatattcaatgttgctgttaaagaatctt-3'<br>aagattctttaacagcaacattgaatataattaaaccaaatagaatcttgttaacaaagtcag-3' |
